# Exploring the bistable equilibrium of methylated CpG DNA recognition by the MBD2 protein

**DOI:** 10.1101/2025.06.30.662303

**Authors:** Senta Volkenandt, Julia Belyaeva, Torry Li, Matthias Elgeti, Ralf Metzler, Petra Imhof, David C. Williams, Mahdi Bagherpoor Helabad

## Abstract

Methyl-CpG binding domain 2 (MBD2) is a critical epigenetic regulator that selectively binds methylated CpG dinucleotides, key marks controlling gene regulation and chromatin organization. Understanding the interactions and conformational dynamics underlying this high selectivity is essential to elucidate MBD2’s regulatory role. Here, using extensive classical MD simulations totaling over 277 *µ*s, we explored the formation of the MBD2–mCpG recognition complex. By initially positioning MBD2 one base pair downstream of its target, we observed its transition to the target site within microseconds. Notably, upon binding, MBD2 adopts two distinct stable conformations: a primary state closely resembling the X-ray crystal structure, and a secondary state of reduced affinity that nevertheless retains comparable selectivity for mCpG. Our results establish S189 as a key macro-switch; loss of its interaction with the methylcytosine backbone shifts the equilibrium toward the secondary state. This is corroborated by MD simulations of the S189A mutant, which preferentially adopts the secondary state-like conformation. Complementary NMR experiments confirm that S189A mutation does not alter mCpG selectivity, while fluorescence polarization measurements reveals a reduced binding affinity, consistent with our MD simulations results. Together, these findings indicate that MBD2 binding to methylated CpG involves a bistable equilibrium, providing new insights into how high affinity and adaptability are balanced in epigenetic recognition. In a broader context, our findings suggest that such alternative bound-state equilibria may represent an inherent feature of specific protein–DNA complexes.

## 1 Introduction

In vertebrates, CpG dinucleotides—cytosines preceding guanines—are almost exclusively the sites of cytosine methylation, commonly represented as mCpG. In humans, approximately 75% of CpG sites are methylated.^1,2^ In cells, 5-methylcytosine (5mC) represents a primary epigenetic modification, where a methyl group is added to the 5-carbon position of the cytosine base.^3,4^ This modification is involved in processes such as aging,^5^ embryonic development and differentiation,^6,7^ nuclear reprogramming,^8^ and tissue-specific gene expression.^9^ The symmetric nature of methylated cytosines in mCpG provides a distinct conformational signal that allows specific domains of some proteins to selectively recognize it.^10^ The main class of proteins that selectively recognize mCpG dinucleotides is the methyl-CpG binding domain (MBD) family. The MBD family, which includes MeCP2, MBD1, MBD2, MBD3, and MBD4, is primarily known for its role in epigenetic regulation and its involvement in signaling events through various protein complexes, with a key function in driving gene repression.^11,12^ The DNA-binding domain, consisting of about 70 residues, is a conserved domain among MBD proteins and exhibits notable structural similarity throughout the family.^13,14^ However, mCpG selectivity depends on the sequence context of individual MBDs. For instance, MBD3, which is highly homologous to MBD2 among MBD proteins, contains a phenylalanine (F34 in MBD3) instead of the highly conserved tyrosine (Y178 in MBD2) found in other MBD family members, resulting in significantly reduced selectivity and binding affinity for mCpG.^15,16^ A pair of conserved arginine residues (R166 and R188 in MBD2), known as arginine fingers, is essential in MBD family members for forming base-specific hydrogen bonds with the symmetric guanine residues within the mCpG dinucleotide and for engaging in cation–*π* interactions with the corresponding methylcytosines.^14,17^

Among MBD proteins, MBD2 exhibits the highest affinity and selectivity for mCpG sites over non-methylated CpG sites (up to 100-fold).^18–21^ The function of MBD2 has been linked to various conditions, including cancer, immune system dysfunction, and neurodevelopmental disorders.^2,22–24^ In cells, MBD2 is an essential component of the nucleosome remodeling and deacetylase (NuRD) complex, which is responsible for chromatin modification and gene silencing by directing the NuRD complex to mCpG islands.^25,26^ For instance, the role of the MBD2–NuRD complex and DNA methylation in silencing the fetal *γ* globin gene in human erythroid cell models is well established.^27,28^ Moreover, MBD2, through its function in chromatin modification, strongly mediates the transition of chromatin from an open to a compacted state.^26^ Studies have shown that MBD2 can engage mCpG sites across a range of DNA conformations—from linear DNA to bent or nucleosome-bound configurations.^17,21,29^ Therefore, an important question is how MBD2 adjusts its high-affinity binding to specific DNA target sites in response to variations in DNA geometry.

MBD2’s function is closely tied to its structural properties and precise mCpG recognition. In the MBD2– mCpG specific complex, in addition to base-specific interactions involving R166, R188, and Y178, there are other residues in MBD2 that establish hydrogen bonds with the DNA backbone.^14,30^ For instance, residue S189, located in the loop connecting *β*3 and *α*1 (Figure 1A), forms a direct hydrogen bond with the backbone of methylcytosine. Mutational studies on this residue in other MBD proteins, such as MBD1 (S45A) and MeCP2 (S134A), show that this mutation reduces their affinity for mCpG by approximately half^31,32^ and likely leads to altered protein conformation relative to DNA, due to the loss of interactions between the protein loop region and the DNA backbone. Importantly, the S134C mutation in MeCP2 has been shown to be pathogenic, causing Rett syndrome; similarly to S134A, S134C reduces the protein’s binding affinity for mCpG—consistent with a loss of function.^32–34^.

**Figure 1.**
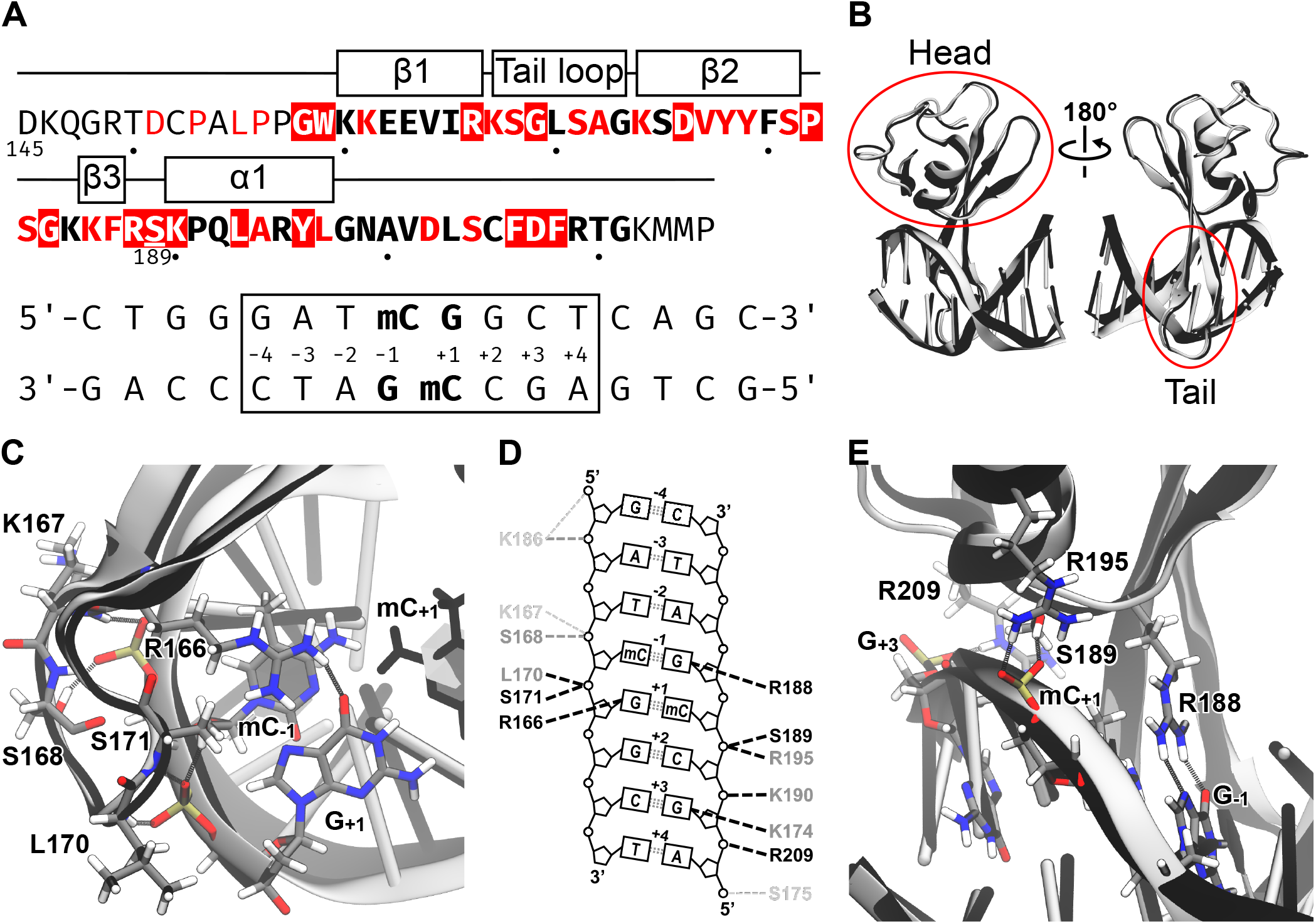
**A** Top: Amino acid sequence of MBD2 with the secondary structure indicated on top; highly conserved residues among the MBD proteins are indicated by a red background, mostly conserved residues are written in red and the core sequence for all analyses is bold. Additionally, S189 is highlighted. Dots help in identifying the residue numbers by marking the tens place. Bottom: DNA sequence of the recognition and interrogation simulations with numbering scheme and the mCpG dinucleotide highlighted in bold. Analyzed eight central basepairs are highlighted by a box. **B** Medoid structure (black) and reference model (gray) of the recognition simulations show high structural similarity with an RMSD of 1.35 Å. The head and tail domain of MBD2 are indicated. **C** Cartoon representation of the medoid structure’s tail domain of MBD2 bound to DNA (black), with hydrogen bond interactions indicated. For comparison, the reference model is shown in gray. **D** Hydrogen bond interactions between MBD2 and DNA observed in the recognition simulations during the 300–1000 ns interval. The color of each residue indicates the occupancy (strength) of its corresponding hydrogen bond: light gray for [0.25, 0.50) (weak), dark gray for [0.50, 0.75) (moderate, and black for [0.75, 1.00] (strong). The thickness and color of each connecting line represent the relative error: black for [0.00, 0.25], dark gray for (0.25, 0.50], and light gray for (0.50, 0.75]; otherwise, a hydrogen bond is not shown. A square bracket indicates the inclusion of a value, while a round bracket indicates the exclusion, so that the intervals do not overlap. **E** Cartoon representation of the medoid structure’s head domain of MBD2 bound to DNA (black), with hydrogen bond interactions indicated. For comparison, the reference model is shown in gray.

Studies of MBD2 dynamics on DNA have shown that methylation significantly restricts MBD2 diffusion, whereas on non-methylated DNA, it diffuses more rapidly.^21^ It has been shown that MBD2 remains bound to mCpG islands and cannot escape for timescales ranging from seconds to minutes.^35^ Nevertheless, MBD2 rapidly exchanges between closely spaced mCpG sites on a sub-millisecond timescale, therefore showing flexibility of MBD2 bound to mCpG.^21,36^ For instance, decreasing the mCpG separation from twelve to eight base pairs accelerates MBD2 exchange between the two mCpG sites, shifting it from an intermediate to a fast NMR timescale.^35^ In such a short distance between mCpG sites, sliding is the most likely motion enabling such efficient exchange between the two mCpG sites. In the target search process of DNA-binding proteins, three key driving factors are: (1) the protein’s exploration of nonspecific sites, primarily via sliding and hopping^37^; (2) the transition from nonspecific search to specific recognition; and (3) the formation of the specific bound state.

Consistent with experimental studies on the kinetics of the target search by DNA-binding proteins,^38–43^ computational studies have characterized the sliding process on different DNA-binding proteins at atomic and coarse-grained levels.^37,44–55^ However, at the atomic level, the search-to-recognition transition and the formation of the specific complex remain poorly understood. Understanding the molecular-level dynamics of such transitions, by using molecular dynamics (MD) simulations (now feasible at microsecond time scales thanks to powerful computational tools), can help predict specific protein–DNA complex structures that are difficult to resolve experimentally.

Therefore, elucidating the underlying dynamics and binding events governing MBD2’s recognition of mCpG sites is crucial for understanding its role in epigenetic regulation.

Here, using extensive MD simulations (in total, 277 *µ*s, Table S1), we investigated the recognition of the mCpG dinucleotide by MBD2. Our results reveal that MBD2 adopts two stable binding states with mCpG DNA, consisting of a primary state equivalent to the X-ray-resolved structure and an alternative stable secondary state. The MD results, validated by NMR experiments, reveal that the key residue distinguishing these states is S189, whose interaction with or detachment from the methylated cytosine’s backbone directs the MBD2–mCpG complex toward the primary or secondary state, respectively. We suggest that the bistable binding behavior at the mCpG site may serve as a mechanism by which MBD2 optimizes the balance between binding specificity and dynamic adaptability.

### 2 Material & Methods

### 2.1 Molecular dynamics simulation

#### Specific protein-DNA complexes

The initial specific protein–mCpG DNA complex was based on atomic resolution coordinates adapted from an NMR structure (PDB ID: 2KY8^30^) and an X-ray crystal structure (PDB ID: 7MWK) of the MBD2 protein. In the NMR structure, two EDTA–thymidine residues that are used for paramagnetic resonance enhancement purposes are mutated to standard thymine residues. The flanking sequence to the mCpG in the X-ray structure was also mutated to that of the NMR structure sequence for consistency. Therefore the DNA sequence used in our simulations is 5′-CTGGGAT<u>**mCpG**</u>GCTCAGC-3′. Two terminal base pairs are also added in the form of a B-DNA model to the DNA sequence to diminish the terminal effects on the central protein–DNA complex. Simulations of the S189A mutation were conducted starting from the X-ray structure. The central eight nucleotides are indexed as G_*−*4_A_*−*3_T_*−*2_**mC**_*−*1_**G**_+1_G_+2_C_+3_T_+4_ with the same index numbers in its complementary strand. Starting from the constructed MBD2–DNA specific complex, we performed eight independent 1-*µ*s MD replicas for each system: the X-ray-derived complex (MD runs R*i, i* = 1,…, 8), the NMR-derived complex (MD runs I*i, i* = 1,…, 8) and the X-ray-derived S189A mutant complex (MD runs S189A_*i*_, *i* = 1,…, 8). Independent replicas were initiated from equilibrated configurations, with initial velocities randomly assigned from the Maxwell–Boltzmann distribution (see Table S1).

#### Nonspecific protein-DNA complexes

We constructed two 18 bps B-DNA molecules with the sequences **DNA 1**: 5′-TCCTGGGAT**mCpG**GCT CAGCC-3′ and **DNA 2**: 5′-AGGGGCmCGG**mCpG**GCTGGCTA-3′ using Chimera.^56^ After aligning the central six AT**mCpG**GC base pairs of this B-DNA model with the equivalent region in the NMR structure, we introduced mutations to mimic the effect of 1 bp shifting the DNA sequence downstream, thereby mimicking the condition where the MBD2 protein is located one base pair upstream from the mCpG site. With the MBD2–DNA nonspecific complex constructed, we conducted seven independent replicas with DNA 1 (named S*i*; *i* = {1,…, 7}) and nine independent replicas with DNA 2 (named S_*N*_ *i*;*i* = {1,…, 9}) of long unbiased MD simulations of ~224 *µ*s in total (see Table S1). Independent replicas were initiated from equilibrated configurations, with initial velocities randomly assigned from the Maxwell–Boltzmann distribution.

#### MD simulations protocol

All molecular dynamics simulations were performed with the GROMACS 2018.4 and GROMACS 2022.5 package^57^, using AMBER14SB^58^ and AMBER parmbsc1^59^ force fields for protein and DNA, respectively. For the methylated cytosine (5mC), we used the combined protocol suggested by K. Liebl et al.^60^ in order to increase the compatibility of the whole DNA force field. For the partial charges of the mCpG base atoms we used the parameters by A. Carvalho.^61^ The systems were solvated with TIP3P water^62^ and potassium (K+) ions were placed randomly to neutralize the system. We also added a salt concentration of 150 mM KCl to mimic physiological conditions. All simulations were initially energy minimized for 5 000 steps using the steepest descent algorithm, followed by 100 ps MD simulation in the NVT ensemble with positional restraints of 1 000 kJ mol^*−*1^ nm^*−*2^ on all heavy atoms of protein and DNA.

Subsequently, positional restraints were gradually lowered to 100 kJ mol^*−*1^ nm^*−*2^ in three 100 ps steps, followed by MD production simulations without positional restraints in the NPT ensemble with a 2 fs time step and trajectory output each 20 ps. In order to preserve a base-paired geometry consistent with B-DNA and prevent melting and terminal fraying, we applied 1 000 kJ mol^*−*1^ nm^*−*2^ distance restraints between hydrogen bond partners and the N1/N9 atoms of the terminal bases on opposing strands. The v-rescale^63^ and Parrinello-Rahman^64^ algorithms were used to keep the temperature and pressure at 300 K and 1 bar, respectively. The van der Waals interactions were switched off between 1.1 and 1.2 nm and long-range electrostatics were treated with particle-mesh Ewald^65,66^ and periodic boundary conditions. The covalent bonds of hydrogen atoms were constrained during the simulations via the LINCS algorithm.^67^

#### Biased MD simulations

##### Setup of the funnel potential

To prevent artificial sliding of the protein along the DNA during steered molecular dynamics (SMD) simulations, we applied a funnel potential using the FUNNEL PS module in PLUMED, following the official tutorial guidelines.^68,69^ A schematic of the funnel potential configuration is shown in Figure S25A. Defining funnel potential in PLUMED we work with two components of the system: the macromolecule (here, DNA) and the ligand (here, MBD2 protein). The macromolecule serves as a reference for aligning trajectory frames and reconstructing the guiding axis of the funnel during simulation (Figure S25B–C). The reference structure included the polar heavy atoms of the central mCpG base pairs, as well as those from three adjacent base pairs upstream and downstream. The ligand can be specified as the center of mass (COM) of a group of atoms. We defined it as the COM of C*α* atoms of the MBD2 protein, excluding N- and C-terminal residues. The direction of the funnel was defined by a line connecting two user-defined points, A and B. In our setup, this axis was aligned with the vector between the centers of mass (COMs) of the DNA and the MBD2 protein in the initial frame of each simulation. Parameters MINS and MAXS determine the position of the lower and upper walls of the funnel that are perpendicular to the A/B guiding line of the funnel. The Zcc parameters determine a point, where the cylindrical part transforms to the cone part of the funnel. Parameters MINS, MAXS and Zcc were set individually for each of three systems via manual inspection using PyMol, version 2.5 Schrödinger, LLC.^70^ For all systems, the funnel radius (Rcyl) was set to 0.1 nm, and the cone angle (*α*) was set to 10°. These values were chosen to allow sufficient freedom for protein rotation and translation during unbinding, while still limiting unphysical lateral movement along the DNA axis.

##### Steered molecular dynamics

SMD simulations were performed using representative structural models of the WT-primary, WT-secondary, and S189A MBD2–DNA complexes.^71^ The system setup and force field parameters were consistent with those used in the long-timescale unbiased molecular dynamics (MD) simulations. For each system, 20 independent SMD simulations were run, each lasting 80 ns. Simulations were performed using GROMACS, version 2023 patched with the PLUMED, version 2.9.0.^57,68,72,73^ The pulling coordinate was defined as the distance between the centers of mass (COM) of selected protein and DNA atoms (denoted as “d com”). The protein COM was calculated using C*α* atoms, excluding flexible N- and C-terminal residues, while the DNA COM was computed using the same atoms included in the alignment reference for the funnel potential. The maximum force constant during pulling was set to 2500 kJ mol^*−*1^ nm^*−*2^ for WT-primary state and 2000 kJ mol^*−*1^ nm^*−*2^ for both WT-secondary state and S189A. A higher force constant was required for WT-primary due to its stronger binding affinity to DNA. We would like to emphasize that SMD simulations were not used for direct binding free energy calculations but rather to generate smooth unbinding trajectories. The restraint force was initially set to zero and gradually increased to its maximum value over the first 0.5 ns. At 0.5 ns, a harmonic potential with the maximal force constant was applied at d com = 1.6 nm to stabilize the system before initiating the pulling. The protein was then gradually pulled away from the DNA, increasing d com to 4.3 nm over the course of 80 ns under constant force. The unbinding trajectories of MBD2 from DNA for each of the three systems were selected in two steps. In the first step, we plotted the reaction coordinate (d com) and the value of the acting funnel potential over time. For each system, we selected three trajectories in which d com increased steadily while the funnel potential remained relatively low. This approach minimized the likelihood of artifacts caused by the funnel potential and ensured that protein unbinding proceeded smoothly. In the second step, we visually inspected these selected trajectories using PyMOL. For each system, we chose one trajectory where the protein exhibited minimal or no rapid rotational movements during unbinding. These final three trajectories were used as the basis for Umbrella Sampling simulations.

##### Umbrella sampling

Umbrella sampling simulations were performed for three MBD2–DNA systems: WT-primary, WT-secondary, and S189A. For each system, a single steered molecular dynamics (SMD) trajectory obtained in a previous step was used. From each SMD trajectory, 35 frames were selected along the unbinding pathway. The first frame corresponded to the fully bound MBD2–DNA state, while the final frame was defined as the point at which all MBD2–DNA contacts were lost. The remaining frames were uniformly distributed between these two endpoints. Each selected frame served as the starting structure for an independent umbrella sampling window. Each window was simulated for 80 ns, resulting in 35 umbrella sampling simulations per system.^68,74^ This simulation length and the number of sampling windows were sufficient to ensure satisfactory histogram overlap (Figure S27). Because biasing potentials were applied in each umbrella sampling simulation, free energy profiles were reconstructed using the binless WHAM reweighting method (all scripts are provided in the Zenodo repository). Bias contributions from all 35 umbrella sampling windows were included in the reweighting procedure. To assess statistical uncertainty, the 80 ns trajectory from each window was divided into 16 consecutive 5-ns intervals (0–5 ns, 5–10 ns, …, 75–80 ns). For each interval, a separate free energy profile was reconstructed by reweighting the corresponding time segment across all 35 independent umbrella sampling replicas. The first 30 ns of data (intervals 0–5 ns through 25–30 ns) were discarded as equilibration.^75,76^ The median profile and standard error of the mean (SEM) were calculated from the remaining ten intervals (30–35 ns through 75–80 ns) to obtain the final free energy estimate and its uncertainty (Figure 5E). Convergence of the free energy profiles was evaluated by reconstructing profiles using cumulative trajectory data of increasing length: 0–10 ns, 0–20 ns, …, 0–80 ns. To illustrate convergence in the second half of the simulation, profiles corresponding to 0–40 ns, 0–50 ns, 0–60 ns, 0–70 ns, and 0–80 ns are shown in Figure S28.

### 2.2 Experiment

#### Protein expression and purification

We expressed and purified HsMBD2MBD(S189A) as described previously,^77^ with only minor changes. In brief, the isolated MBD(S189A) domain (amino acids 150–214) was cloned into a modified pET32a vector^78^ with a TEV protease site immediately preceding the domain. We transformed the resulting plasmid into the Rosetta II (DE3) (Invitrogen) *E. coli* strain, which we grew and induced expression as previously. We purified the expressed fusion protein by nickel affinity chromatography, removed the thioredoxin N-terminal tag by TEV protease cleavage, and further purified the domain by gel filtration chromatography (Superdex-200 26/60, Cytiva).

#### DNA purification

We purchased complementary 17-base pair oligonucleotides (5′-GAGGCGCT(mC)GGCGGCAG-3′) with a single symmetrically methylated site from Integrated DNA Technologies (IDT). The DNA was annealed and purified by ion exchange chromatography (Source 15Q, Cytiva). We purchased a 3′ FAM labeled forward oligonucleotide for fluorescence polarization measurements.

#### Fluorescence polarization (FP) analysis

We buffer-exchanged protein and DNA samples into 10 mM HEPES pH 7.5, 50 mM NaCl, 3 mM MgCl2, 0.1 mM EDTA, 1 mM DTT, and 0.02% sodium azide. The protein was serially diluted in the presence of 10 nM 3’FAM-labeled DNA in a 96-well plate, and fluorescence polarization was measured on a CLARIOstar microplate reader (BMG Labtech). We fit the resulting data to a general equation for two-state binding

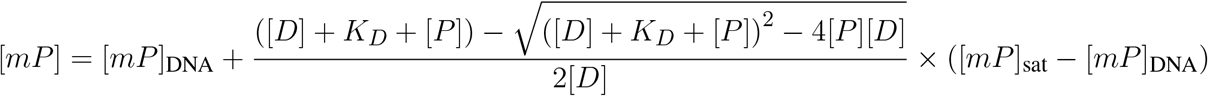

with Prism graphing software (GraphPad). [D] and [P] are the DNA and protein concentration, respectively; [mP]DNA and [mP]sat are the polarization for free and protein saturated DNA, respectively. The serial dilutions and measurements were performed in triplicate, and the experiment was repeated three times.

#### Nuclear magnetic resonance spectroscopy

^15^N-labeled protein bound to the methylated double-stranded DNA was buffered exchanged into 10 mM NaPO4, pH 6.5, 0.02% sodium azide, 1 mM dithiothreitol, and 10% ^2^H_2_O and concentrated to approximately 0.3 mM. NMR spectra were collected on a Bruker Avance III 850 MHz instrument at 25 °C, and data were processed and analyzed with NMRPipe^79^ and CcpNmr^80^, respectively.

### 2.3 Analyses

#### Native Contact Analysis

Native versus non-native interactions between protein and DNA are investigated using the native contact concept, introduced by Robert B. Best et al.^81^ The fraction of native contacts *Q* can be computed as

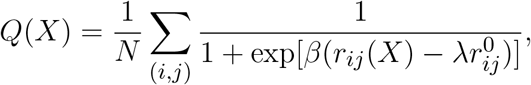

where the summation is over all *N* pairs of native contacts (*i, j*), and *r*_*ij*_(*X*) and 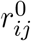 are the distances between atoms *i* and *j* in configuration *X* and in the reference conformation, respectively. As suggested in^81^ we used a smooth parameter *β* of 5 Å ^*−*1^ and a fluctuation factor *λ* of 1.8.

#### Similarity index

To measure the structural similarity between the search and recognition states, we adopted the protocol of Leven et al.^44^, where the parameter *χ*_*i*_ is calculated for each protein residue forming specific interactions with DNA in the bound complex as

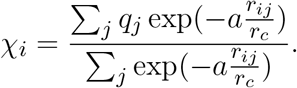

In this definition, *j* corresponds to all protein residues within a cutoff radius of *r*_*c*_ = 8 Å from residue *i. r*_*ij*_ denotes the distance between residues *i* and *j, q*_*j*_ represents the point charge associated with residue *j*, and *a* = 5 is the exponential factor. The total similarity index *χ* is defined as the mean of all residue-specific *χ*_*i*_ values and spans from − 1 to +1, with higher *χ* values corresponding to lower frustration and a higher degree of similarity between the search and recognition states.

#### RMSD clustering

The RMSD clustering was performed with the gromos^82^ method of GROMACS^57^ module `gmx cluster` using each 50th frame and a cut-off of 0.25 nm for WT simulations, and each tenth frame and a cut-off of 0.22 nm for the S189A mutation simulations. Cluster centers, meaning representative structures, are the structures with the minimum average RMSD to all other structures within the respective clusters. Skipped frames have been assigned to the cluster of the frame closest in RMSD taken only into account clusters with at least 0.5 % of all data points. More details about the choice of thresholds can be found in the Supplementary Information.

#### Principal component analysis (PCA)

Principal component analysis has been performed using GROMACS.^57^ With gmx covar we first calculated the mass-weighted covariance matrix and its eigenvalues and eigenvectors for the fluctuations of C_*α*_ (for protein) and P (for DNA) atoms. Then the trajectory is projected onto these eigenvectors using `gmx aneig`. The cosine content of a principal component was calculated with `gmx analyze` to ensure that it does not represent random diffusion.^83^

#### DNA parameters

The DNA conformation, including the geometrical inter and intra bp parameters, DNA grooves, and backbone dihedral angles were calculated with Curves+/Canal^84,85^. The DNA backbone BI/BII substates are determined by the *ε* and *ζ* dihedral angles, where *ε − ζ <* 0 means BI configuration and otherwise BII.

#### Hydrogen bonds

Hydrogen bonds have been determined based on geometric criteria. The angle between hydrogen, donor and acceptor atom of a hydrogen bond deviated no more than 42° from linearity and the distance between hydrogen and acceptor atom has to be lower than 2.2 Å.

#### Contact area

The contact area between protein and DNA has been determined based on the solvent accessible surface are (SASA)^86^ of the protein (*A*_prot_), the DNA (*A*_DNA_) and the protein-DNA complex (*A*_complex_) as

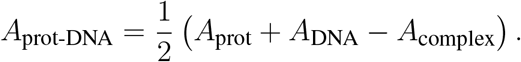

## 3 Results

### 3.1 The specific complex exhibits high stability

The highest specificity and affinity of MBD2 for mCpG, compared to other MBD proteins, suggests the formation of a highly stable MBD2–mCpG complex. To evaluate the stability of the MBD2 bound specifically to mCpG, we carried out eight 1 *µ*s MD simulations (R1–R8) starting from the X-ray structure (PDB ID: 7MWK), named *recognition simulations*. Figure 1A illustrates the MBD2 protein sequence along with the DNA sequence containing mCpG used in this study. RMSD analysis of the recognition simulations reveals that the MBD2–mCpG complex maintains a stable conformation with an average RMSD of 1.87 *±* 0.35 Å (for full time series see Figure S1**A**). Figure 1B depicts the cartoon model alignment of the medoid structure calculated from all recognition simulations with respect to the MD-equilibrated model based on the crystal structure (hereafter, ‘reference model’), demonstrating high structural similarity with an RMSD of 1.35 Å between the two structures (see Table S2). Two key subdomains of the MBD2 DNA-binding domain, here referred to as the head and tail domain (Figure 1B), establish unique hydrogen bonds with mCpG DNA. Figures 1C–E illustrate the hydrogen bond interactions and their occupancies calculated from the recognition simulations. In the head domain, the two residues R188 and S189 form strong hydrogen bonds with the base of G_*−*1_ and the backbone of mC_+1_, respectively (Figure 1D,E). The formation of the S189–mC_+1_ hydrogen bond also facilitates the moderate interaction of R195 with mC_+1_. In the tail domain, the key base-specific interaction, illustrated in Figure 1C, is between R166 and G_+1_, the complementary base of the methylated cytosine mC_+1_. Furthermore, tail domain residues L170 and S171 jointly contribute strong hydrogen bonds to the backbone of G_+1_ (Figure 1C,D). This is also supported by the water-mediated interaction of Y178 with mC_*−*1_ (illustrated in Figure S2A,B). Outside of mCpG (Figure 1D), the only moderate base-specific interaction occurs between K174 and G_+3_. Residue R209 also forms a strong hydrogen bond interaction with the backbone of G_+3_. These detailed, strong interactions—clearly maintained throughout our recognition simulations—are crucial for stabilizing the specific MBD2–mCpG complex.

### 3.2 MBD2 being at the target site does not guarantee recognition

The first deposited PDB structure of the MBD2–mCpG complex was resolved using NMR spectroscopy.^30^ However, owing to the low proton density observed in the corresponding NMR study, the DNA in this structure was modeled in part on the B-DNA conformation. As shown in Figure S3, a comparison of the NMR-resolved structures with the X-ray structure reveals differences in the positioning of the head and tail domains as well as the DNA structure relative to the X-ray structure. To assess the influence of these structural variations on the conformational stability of the MBD2–mCpG complex, we performed eight additional MD replicas (I1–I8), initiated from the NMR structure (PDB ID: 2KY8), named *interrogation simulations*. In contrast to the recognition simulations, RMSD analysis of the interrogation simulations indicates lower stability, with higher RMSD values relative to the reference model (for time series see Figure S1B). However, three out of the eight interrogation MD replicas, i.e. I2, I3, and I7 converged to the reference model based on the X-ray structure (Figure S1B, while the remaining simulations failed to achieve the specific conformation within the 1 *µ*s simulation time span and also remained far from the initial NMR structure, indicating that they were still in a search state (Figure S1B,C). These simulations indicate that mere localization at the target site is not sufficient for formation of the recognition complex; rather, a final transition occurring on a timescale exceeding 1 *µ*s is required to reach the full recognition state.

### 3.3 Fast target localization precedes slow recognition complex formation

After confirming the stability of the MBD2–mCpG recognition complex in recognition simulations and showing that this state is not readily reached from the NMR-based starting structure, we probed target-site search by positioning MBD2 one base pair downstream from its recognition site. This setup enabled us to assess target localization, specific mCpG binding, and the molecular determinants of the transition. We therefore generated a non-native initial complex (Figure S4A) and performed long-timescale MD simulations (seven replicas; see Methods), hereafter referred to as *search simulations* (S1–S7). Initially, we utilized the similarity index (*χ*), introduced by Leven *et al*.^44^ (see Methods), to quantify the difficulty for a DNA-binding protein in transitioning from its search mode to the recognition mode. The similarity index *χ* reflects the overlap between the search and recognition states, where higher *χ* values correspond to lower frustration, stronger similarity, and a reduced free energy barrier, whereas lower *χ* values indicate the opposite. For the MBD2–mCpG complex, our analysis yielded *χ* = 0.35, indicating rather similar MBD2’s conformation between these states. Our search simulations reveal that the transition of MBD2 from its initial configuration to a position localized on the mCpG dinucleotide occurs rapidly within a sub-microsecond timescale, as illustrated in Figure 2A by the RMSD evolution from the initial structure, which is also supported by the rotation angle evolution of MBD2 around the DNA (Figure S5). This rapid sliding motion, however, may also be attributed to a DNA conformational shift from the canonical B-DNA form to a protein-induced adapted conformation. Figure 2B shows the RMSD profiles from the search simulations with respect to the reference model. Notably, four search simulations (Figure 2B, replicas S1, S2, S6, and S7) successfully adopt a structure in agreement with the recognition state, characterized by an RMSD of approximately 2 Å. The observed rotational angle Δ*θ*_1_ ≈ 30° (Figure S5) suggests an approximately 1 bp rotational movement along the DNA in these simulations. In contrast to the rapid sliding motion, the transition from the localized state to the native recognition state occurs on a timescale of several microseconds, as indicated by the non-highlighted regions of replicas S1, S2, S6, and S7 in Figure 2B. This suggests that localization of the protein at the specific DNA sequence does not necessarily result in immediate formation of the specific conformation. This is in agreement with the moderate search-to-recognition transition barrier indicated by the similarity index.

**Figure 2.**
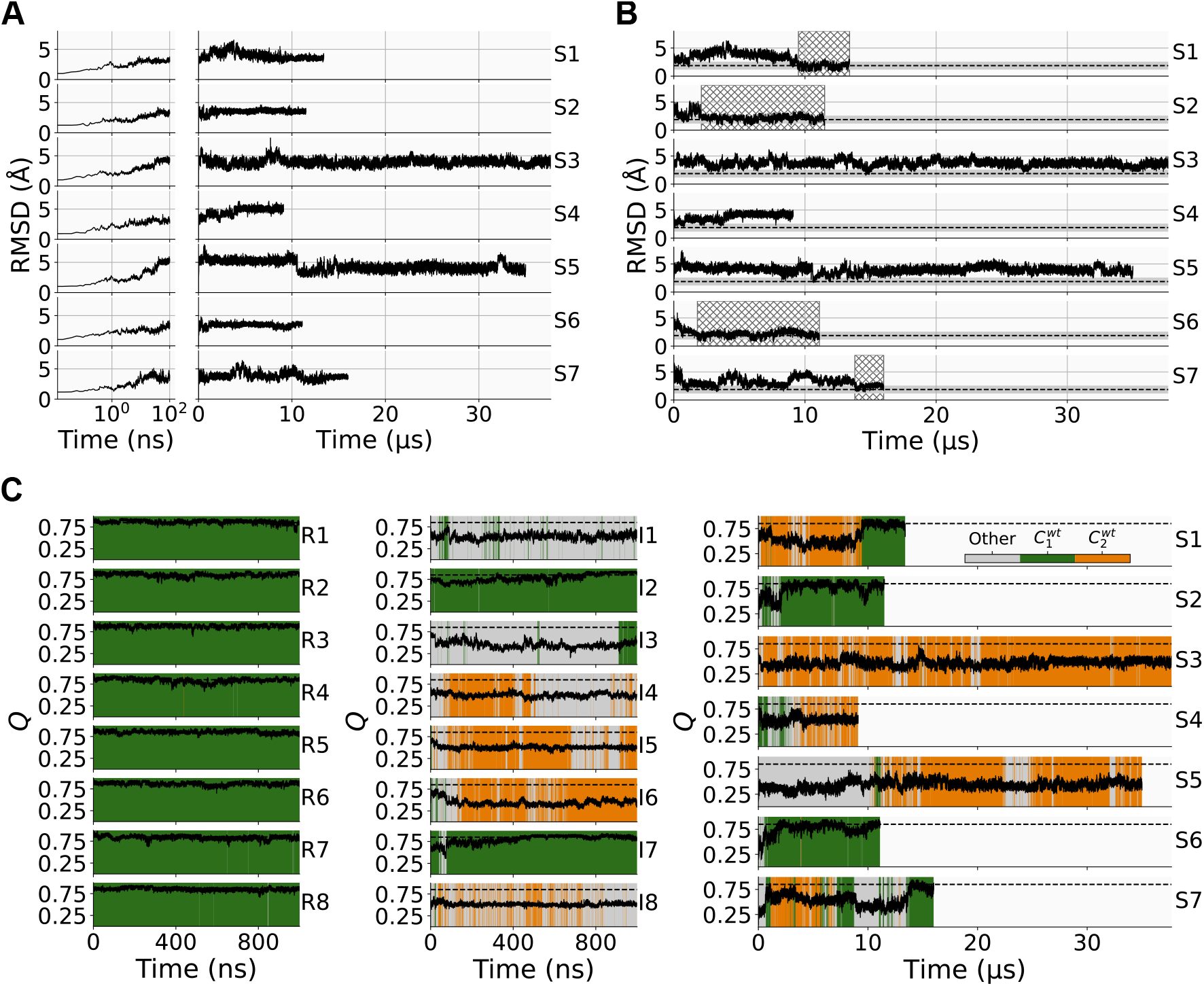
**A** RMSD of the search simulations with respect to the first frame. The first 100 ns are plotted additionally on a log-scale. **B** RMSD of the search simulations with respect to the reference model. The average RMSD of 1.87 Å (300 *−* 1000 ns) of the recognition simulations is highlighted as a dashed line, while the gray area around it additionally indicates the fluctuation around that average as two standard deviations. Parts of the trajectory that are within that range are highlighted by a hatched area. **C** Clustering based on pairwise RMSD with a threshold of 0.25 nm over time for all recognition, interrogation, and search simulations is shown in green, orange and gray for clusters 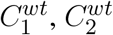 and all others, respectively. The fraction of native contacts *Q* with respect to the reference model over time is shown as solid black line, while the average fraction of native contacts of the specific complex from the recognition simulations is shown as a black dashed line. Note the different time axis scales.

### 3.4 An additional binding state emerges from the search simulations

Despite successful formation of the recognition complex in replicas S1, S2, S6, and S7, simulations S3–S5 fail to reach this state, instead stabilizing at an RMSD of ~4 Å relative to the reference model (Figure 2B). To determine whether the MBD2–mCpG complex in these simulations form distinct, well-defined conformations different from the recognition complex, we performed pairwise RMSD clustering across all trajectories from the search, recognition and interrogation simulations. Additionally, we analyzed the evolution of native contacts (*Q*) in all simulations with respect to the native complex, i.e. the reference model. Figure 2C displays the native contact profiles mapped onto RMSD-based clusters with distinct regions highlighted by different colors.

#### 3.4.1 Cluster analysis reveals two stable conformations

Our cluster analysis identifies two predominant conformational states associated with stable binding: the primary state, represented by cluster 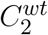, which corresponds to the native recognition complex (depicted in green in Figure 2C), and the secondary state, represented by cluster 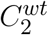, which reflects an alternative binding conformation (depicted in orange). The corresponding pairwise RMSD distribution (Figure S6A) exhibit a valley, supporting separated states. A principal component analysis based on C*α* and P atom fluctuations provides additional support for the separation into these two states (Figure S7). As shown in Figure 2C, *left*, the recognition simulations remain entirely within the primary specific state; with a fraction of native contacts *Q* close to 1 and RMSD clustering showing a single cluster 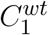, corresponding to the specific complex. However, only two of the interrogation simulations (I2 and I7) achieve a conformation resembling the specific complex (Figure 2C, *middle*). In I3, although the simulation ultimately falls within cluster 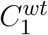, the native contacts are not completely formed, with *Q* ≃ 0.5. This indicates that the specific conformation can only be achieved when both clustering and native contact conditions are fulfilled. The four replicas I4, I5, I6, and I8 in the interrogation simulations adopt conformations closer to cluster 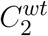. In the search simulations, as noted above, replicas S1, S2, S6, and S7 adopt the primary state, which resembles the specific complex, as indicated in Figure 2C, *right* by their assignment to cluster 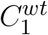 and a native contact value (*Q*) close to one. However, the search replicas S3–S5 predominantly adopt the distinct secondary state, represented by cluster 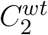 (Figure 2C, *right*). Notably, the secondary state can persist over multiple microseconds (replicas S3 and S5), thus emphasizing its stable nature.

#### 3.4.2 The secondary state adopts a distinct complex conformation

Here, we examine the characteristics that differentiate the secondary state from the primary state. Figure 3A depicts the representative structures of the primary (green) and secondary states (orange) in cartoon illustration with respect to the reference model (gray). The representative primary state closely resembles the reference model (RMSD of 1.8 Å), whereas the secondary state shows a larger deviation (RMSD of 3.6 Å); the two states differ by 4.1 Å (Table S3). The key difference between secondary and primary states (or crystal structure) lies in the distinct positioning of the protein relative to the DNA, primarily due to a shift in MBD2’s head domain accompanied by a minor displacement of the tail domain, resulting in a reconfigured protein–DNA interface. In contrast to the primary state, where one base-pair sliding occurs mainly through pure rotation around a defined axis, sliding in the secondary state involves coupled rotation and conformational changes caused by rearrangement of the MBD2 head domain and DNA. Figures S8 and S9 show two representative examples (S7 and S5) of such sliding along DNA toward the primary and secondary states, respectively. Interestingly, in S5, MBD2 transiently slides even beyond one base-pair before subsequently relaxing back to the stable secondary state (Figures S9).

**Figure 3.**
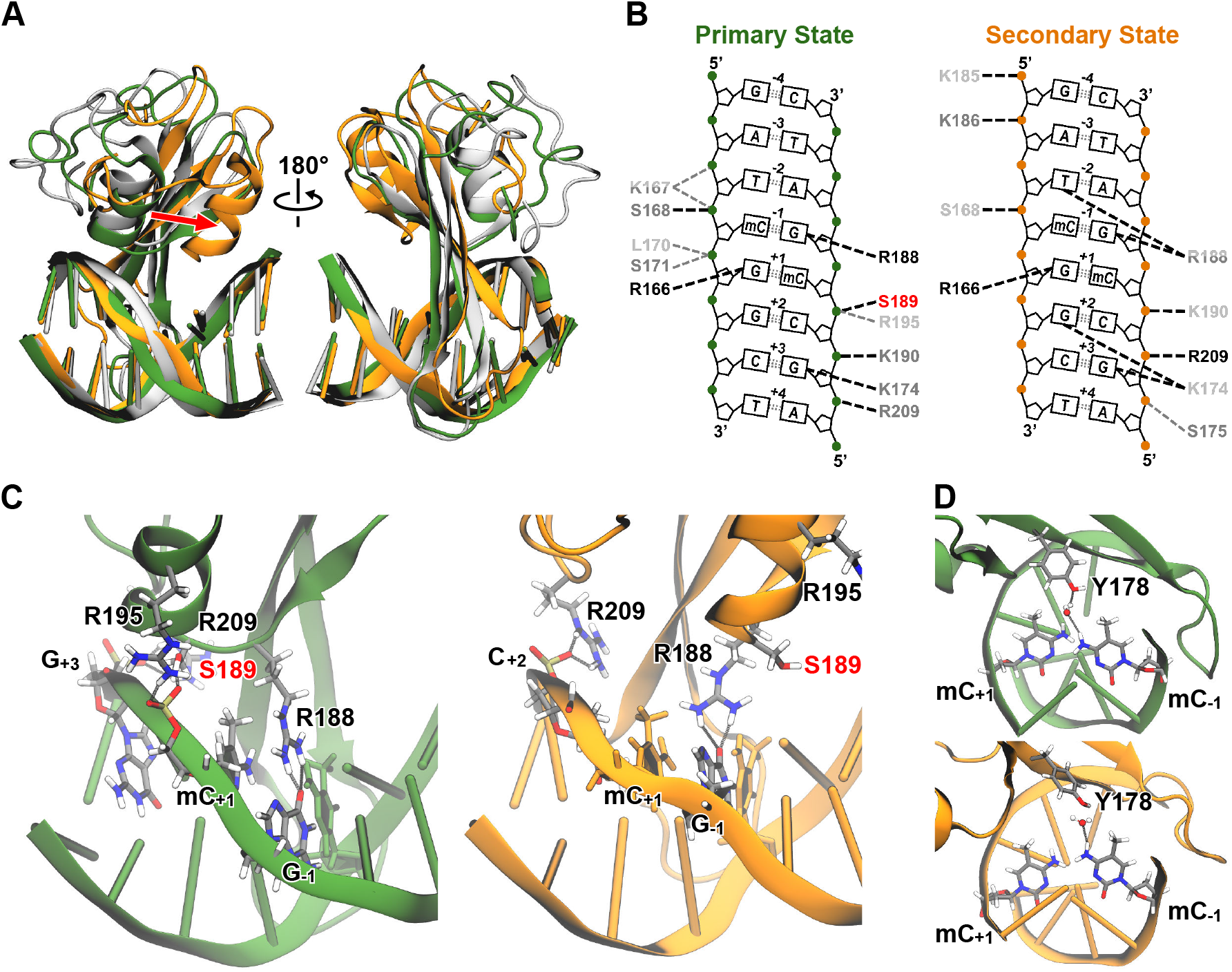
**A** Cartoon representation of the cluster centers (green and orange) corresponding to the clustering shown in Figure 2C and the reference model (gray). A red arrow indicates the shift of the head domain’s helix *α*1 for cluster 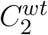 (orange) with respect to the recognition complex. **B** Hydrogen bond interactions between protein and DNA for the two main clusters, which we identify as primary (green) and secondary (orange) states. The color of the residue corresponds to the occupancy (strength) of the corresponding hydrogen bond: light gray for [0.25, 0.50) (weak), dark gray for [0.50, 0.75) (moderate) and black for [0.75, 1.00] (strong). The thickness and color of the line on the other hand represents the relative error of that hydrogen bond: black for [0.00, 0.25], dark gray for (0.25, 0.50] and light gray for (0.50, 0.75]; otherwise, a hydrogen bond is not shown. A square bracket indicates the inclusion of a value, while a round bracket indicates the exclusion, so that the intervals do not overlap. **C** Cartoon representation of the head domain of MBD2 bound to DNA showing the hydrogen bond interactions for the two cluster centers 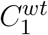 (left, green) and 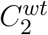 (right, orange), corresponding to the primary and secondary states, respectively. **D** The absence or presence of the water-mediated hydrogen bonds between Y178 and the base of mC _*−* 1_ in clusters 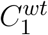 (top, green) and 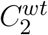 (bottom, orange), respectively.

To examine the difference between these observed bistable equilibrium in more detail, we calculated the hydrogen bond interaction map for the primary and secondary states, as shown in Figure 3B. A comparison of the hydrogen bond interactions of the primary state (Figure 3B, *left*) and those of the recognition simulations (Figure 1D) shows a high degree of similarity. The hydrogen bond maps of the primary and secondary state indicate that the three base-specific interactions crucial for the selective recognition of mCpG DNA—R166, K174, and R188—are also retained in the secondary state. However, a slight shift in hydrogen bond interactions is observed toward the downstream region of the DNA, resulting in altered interactions of residues K174 and R188, which establish additional moderate contacts with G_+2_ and T_*−*2_, respectively. A key loss of interaction contributing to the formation of the secondary state is the disruption of the hydrogen bond between S189 and mC_+1_, which remains strong in the primary state (Figure 3B). The absence of this DNA backbone-mediated interaction is a key factor in the displacement of the MBD2 head domain within the DNA major groove, resulting in formation of the secondary state. Figure 3C depicts the MBD2 head domain in both states, highlighting its conformational shift and the positioning of residue S189 within the DNA-binding interface. As a result of this conformational shift the R209–G_+3_ and K190–C_+2_ hydrogen bonds in the primary state, are rearranged into R209–C_+2_ and K190–mC_+1_ interactions in the secondary state (Figure 3B,C), respectively, meaning they shift one nucleotide downstream. Additionally, the conformational change of the MBD2 tail domain in the secondary state, compared to the primary state, disrupts the moderate interactions of residues K167, L170, and S171, which are observed only in the primary state (Figure 3B and structurally shown in Figure S10A,B). Consequently, contact area analysis reveals a larger buried surface at the protein-DNA interface in the primary state than in the secondary state (Figure S11A), consistent with the stronger hydrogen bond interactions observed in the primary state. The secondary state’s distinct conformation with respect to the primary state also significantly alters Y178’s positioning relative to the mCpG bases. As shown in Figure 3D and Figure S12 (distance distribution), residue Y178 forms a strong water-mediated hydrogen bond interaction with mC_*−*1_ in the primary state, whereas in the secondary state this interaction is largely absent, as supported by the distance distributions between the Y178-OH atom and the N4 atom of mC_*−*1_ and mC_+1_ in Figure S12. The overall MBD2–mCpG contact maps (Figure S13) show distinct contact patterns for the primary and secondary states relative to the recognition simulations, while the primary and recognition conformations differ only subtly. Moreover, our analysis reveals a significant conformational change in the DNA when comparing the primary and secondary state. In particular, we observe a significant change in the backbone BI/BII conformation, a prominent structural parameter that critically influences DNA flexibility and function.^87–93^ As shown for G_*−*1_ in Figure 4A and for mC_*−*1_ and mC_+1_ in Figure S14, the secondary state exhibits distinct, well-defined BI/BII backbone conformations, whereas the primary state remains nearly identical to the specific complex (dashed lines). Figure 4B depicts the backbone of G_*−*1_ in BI and BII conformations, showing that this transition affects not only the base position but also the neighboring mC_+1_, as reflected by the phosphate-group positioning. In addition, the DNA geometric parameters, including inter- and intra-base-pair as well as groove parameters (Figs. S15–S17), reveal considerable conformational changes in the DNA in the secondary state compared to the primary state, whereas the primary state displays a geometry closely resembling to that of the specific recognition complex. This is particularly evident in the mCpG dinucleotide and, to some extent, in the nucleotides directly flanking mCpG, as reflected in the intra-base-pair parameters such as opening, propeller, x–, and y–displacement (Figure S15), as well as inter-base-pair parameters such as shift and twist (Figure S16). Notably, the distinct DNA conformation observed in the secondary state, compared to the primary state, extends not only at the base-pair level but also to the backbone, as evidenced by the minor and major groove parameters (Figure S17).

**Figure 4.**
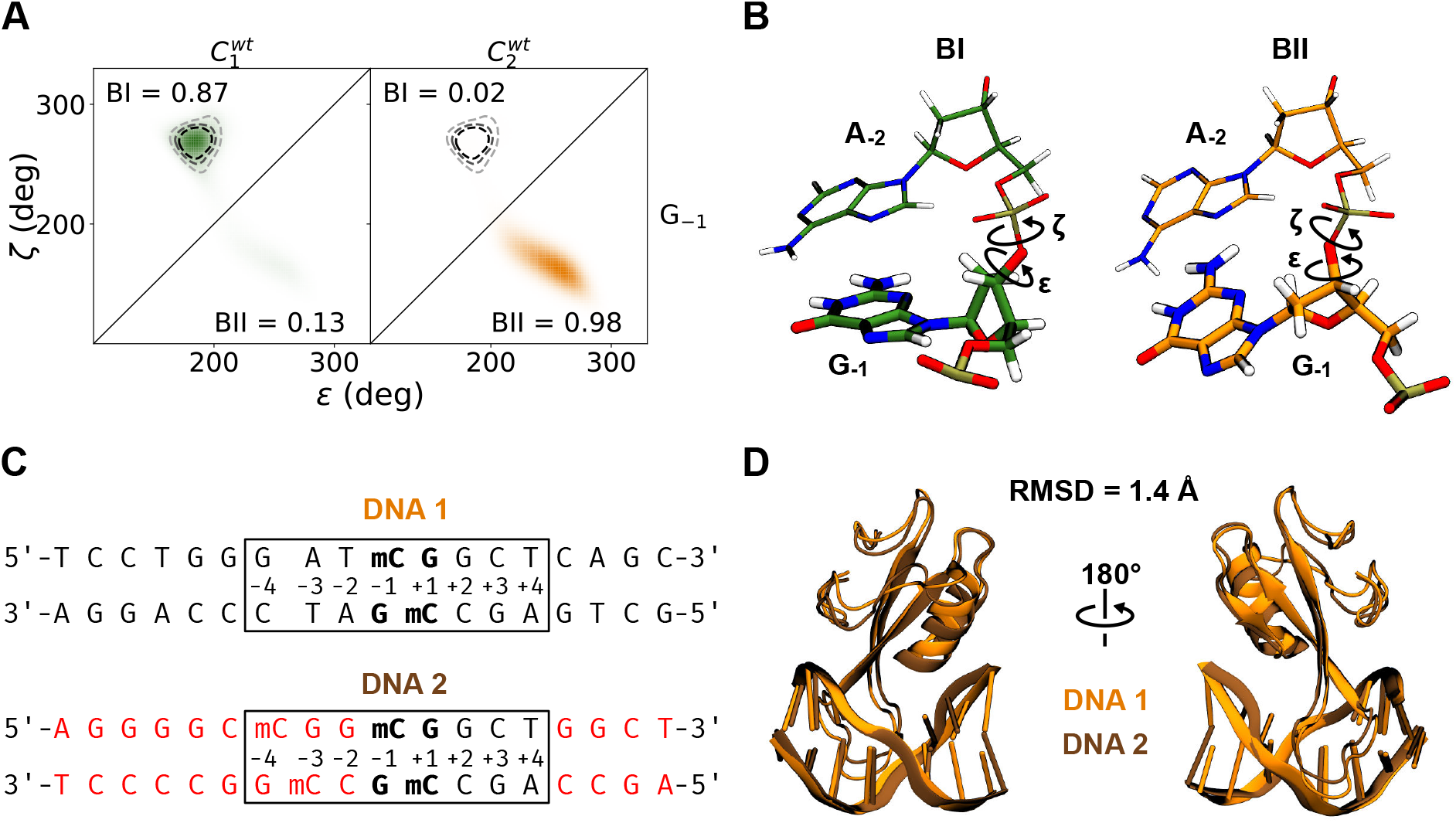
**A** Distribution of backbone angles *ε* and *ζ* representing the backbone conformation, which is either BI or BII, is shown for the guanine G_*−*1_ of clusters 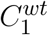 left, green, primary state) and 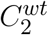 (right, orange, secondary state) as well as for the specific complex (dashed gray lines). **B** Representative structures showing the G _*−* 1_ backbone conformation, which is mainly BI for the primary state (left, green) and mainly BII for the secondary state (right, orange). Backbone angles *ε* and *ζ*, which determine the backbone conformation, are indicated. **C** Comparison of the two DNA sequences used for search simulations, DNA 1 (top) and DNA 2 (bottom). The differences in sequence are shown in red for DNA 2. The central eight basepairs are highlighted by a box and the mCpG dinucleotide in bold. **D** Cartoon representation of the cluster centers for the secondary state of search simulations performed on DNA 1 (orange) and DNA 2 (brown) show high structural agrrement.

To further validate the secondary state as an alternative sequence-specific bound conformation, we performed additional search simulations using a different DNA sequence containing the native mCpG sites at positions −53 and −50 relative to the transcription start site of the *γ*-globin gene promoter in adult erythroid cells, where MBD2 functions as a cofactor in gene silencing.^28^ The new DNA sequence, labeled DNA 2, together with the previously used sequence (DNA 1), is shown in Figure 4C. Similar to the previous search simulations, in the new simulations with DNA 2 (9 replicas, each 10 *µ*s), MBD2 was positioned one base pair downstream relative to the mCpG site at position −50 (Figure 4C, shown in bold). Here, RMSD clustering of the new search simulations (Figure S18A) also reveals two states corresponding to the primary and secondary states observed with DNA 1 (RMSD distances can be found in Table S4), with the S189–mC_+1_ interaction re-emerging as the key macro-switch. Notably, five out of nine new search replicas converge to the secondary state conformation, which closely recapitulates the secondary state observed in the previous simulations with DNA 1, with an RMSD of 1.4, Å between them, as shown by the superimposed structures in Figure 4D (for the primary state see Figure S18B,C.)

Taken together, our results suggest that the secondary state represents an alternative conformational equilibrium distinct from the primary state while preserving the key structural features required for for the formation of a stable, specific MBD2–mCpG complex. Its recurrence in a second, biologically relevant DNA sequence further substantiates this alternative specific bound conformation.

### 3.5 Validating the secondary state via S189A mutation

#### 3.5.1 S189A mutation shifts the MBD2–mCpG complex toward the secondary state conformation

As demonstrated in the previous section, a critical distinction between the primary and secondary state is the absence of the S189–mC_+1_ hydrogen bond interaction in the secondary state. To determine whether the S189A mutation might shift the binding equilibrium toward the secondary state, we performed eight independent 3 *µ*s MD simulations of the specific complex bearing the S189A mutation based on the 7MWK crystal structure (*S189A simulations*). The fraction of native contacts *Q* in the S189A simulations (Figure S19A) reveals that the protein-DNA complex gradually deviates from the native specific configuration over the course of the simulation in nearly all trajectories. Performing RMSD clustering on the entire set of S189A simulations, following the same approach as in the *wild-type (wt)*-simulations, reveals three distinct clusters, as illustrated in Figure S19A. The cluster shown in dark blue (cluster 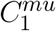) corresponds to a conformation similar to the specific complex (or primary state structure). Figure 5A shows that the cluster center of cluster 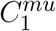 closely resembles the primary state structure (cluster center 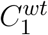, green) with an RMSD of 1.1 Å. It can be mainly found in the beginning of all S189A replicas. Six out of the eight S189A simulation replicas—M1, M3, M4, and M6–M8 (for cluster time series see Figure S19B)—converge toward cluster 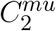) which closely resembles the conformation and binding interactions of the secondary state cluster 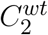 (orange) observed in the *wt*-simulations(Figure 5B), with an RMSD of 2.2 Å This convergence from the primary toward the secondary state is further supported by the native contact fractions of the S189A replicas relative to both states (Figure S19B); specifically, the native contact fraction decreases over time relative to the primary state while increasing relative to the secondary state. Together, this indicates that the protein–DNA contacts in cluster 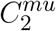 closely align with those observed in the secondary-state complex. The other two replicas, M2 and M5, adopt an intermediate conformation 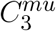, where the head domain positioning on DNA resembles 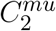, while the tail domain remains similar to cluster 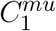, see Figure S20B. However, the DNA structure in 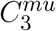 adopts a conformation similar to that of cluster 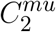, as evidenced by DNA geometry parameters (Figure S21–S23) as well as the BI/BII conformation (Figure S24) Taken together, our simulation results demonstrate that the S189A mutation shifts the MBD2–mCpG conformation toward the secondary state observed in the *wt*-simulations.

**Figure 5.**
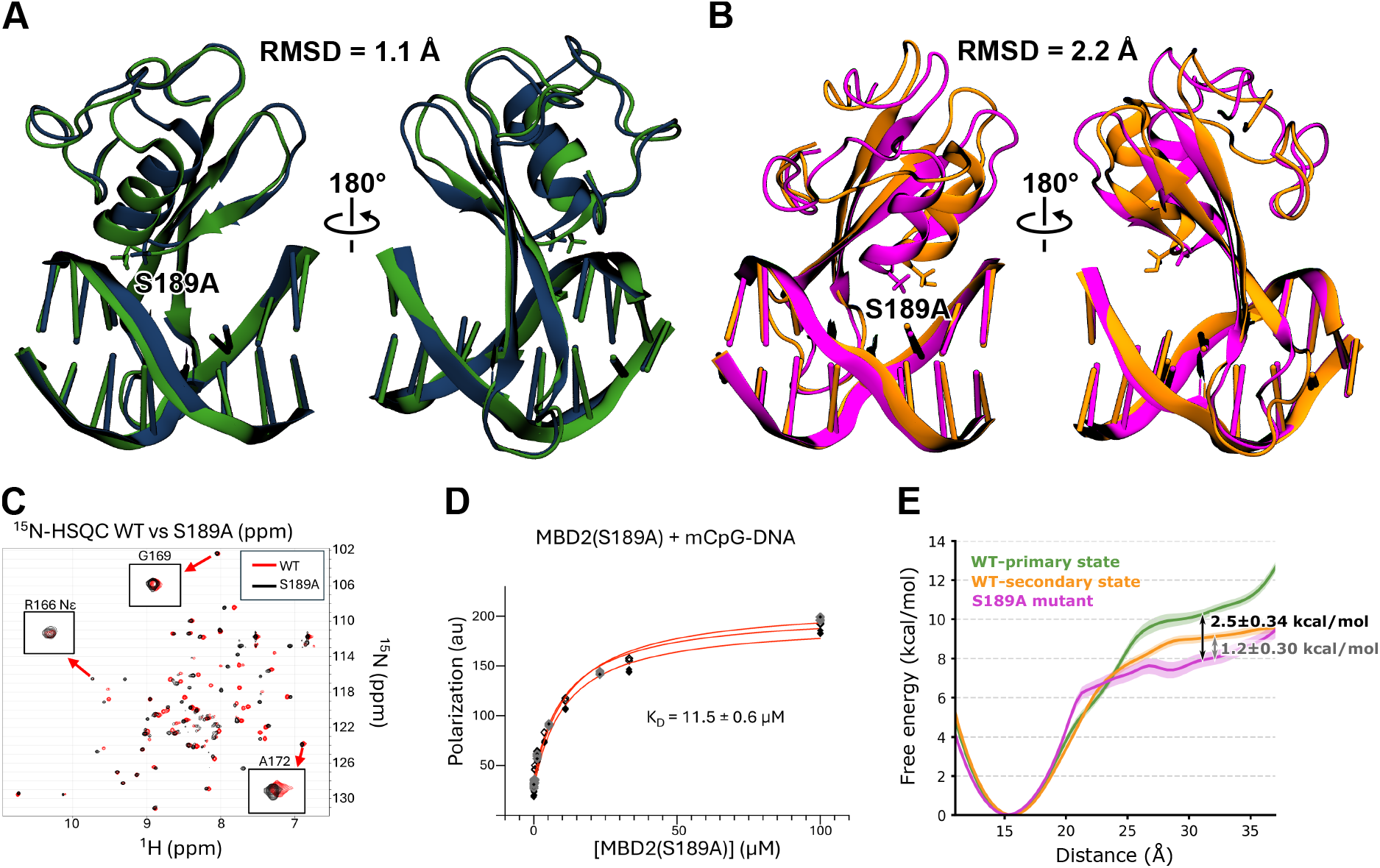
Methylation selectivity of MBD2(S189A). **A** Comparison of cluster center 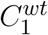 (green) obtained from the wild-type simulations and the cluster center 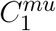 (blue) obtained from the S189A mutation simulations. **B** Comparison of cluster center 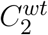 (orange) obtained from the wild-type simulations and cluster center 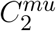 (pink) obtained from the S189A mutation simulations. The mutation is highlighted. **C** An overlay of 2D ^15^N-HSQC spectra is shown for MBD2-MBD wild-type and S189A. Three sub-spectra show expanded views of reporter resonances (R168 N*ϵ*, G169, and A172) that show large chemical shift changes between methylation-specific and nonspecific binding modes. **D** The binding affinity of MBD2-MBD(S189A) for methylated DNA was measured by fluorescence polarization analysis. The fluorescent polarization intensities and corresponding fits are depicted for each of the three replicates (*N* = 3). **E** Binding free energy profiles from Umbrella Sampling simulations for the *wt*-primary state, *wt*-secondary state, and the S189A mutant. Bold lines indicate the statistical mean, and shaded areas represent the standard error of the mean (SEM). Arrows show free energy differences between *wt*-primary and S189A, and between *wt*-secondary and S189A (in kcal/mol).

#### 3.5.2 S189A mutation retains MBD2 selectivity for mCpG but reduces binding affinity

To assess whether the S189A mutation affects MBD2’s selectivity for mCpG, we performed a ^15^N HSQC NMR experiment. Previous studies have identified three cross-peaks in MBD2, corresponding to highly conserved residues G169, R166, and A172, whose chemical shifts reflect mCpG specific binding by MBD2. These three cross-peaks exhibit large chemical shift changes between nonspecific and methylation-specific binding modes.^17,35,94,95^ For example, the cross-peak for ^1^H*ϵ* of R166 shifts over 1 ppm downfield when bound to methylated DNA compared to unmethylated DNA. Our ^15^N HSQC data (Figure 5C) indicate that the cross-peaks corresponding to these residues, i.e. G169, R166, and A172, remain unchanged upon the S189A mutation, demonstrating that this mutation does not alter MBD2’s selectivity for mCpG. In contrast, many of the other resonances show chemical shift changes with the S189A mutation, consistent with a change in the relative orientation between the MBD2 and DNA. Hence, the mutant protein maintains methylation-specific binding even though its orientation with respect to DNA likely changes. Next, to quantify the effect of the S189A mutation on binding affinity, we performed a fluorescence polarization (FP) experiment measuring MBD2–S189A binding to mCpG DNA. As shown in Figure 5D, we measured a dissociation constant of K_D_ = (11.5 *±* 0.6) *µ*M, indicating a ten-fold reduction in binding affinity compared to *wt*-MBD2, which has a dissociation constant of K_D_ = (0.72 *±* 0.06) *µ*M as we have previously determined using the same DNA and experimental conditions.^77^ Therefore, the binding free energy difference as a result of the S189A mutation is (1.67 *±* 0.51) kcal/mol, consistent with the expected energetic contribution of a single strong hydrogen bond interaction.^96,97^ We further sought to evaluate the binding free energy of the MBD2–mCpG complex for both *wt* (primary and secondary states) and S189A variants using advanced enhanced sampling simulation techniques (see Methods). Using steered molecular dynamics (SMD) with a funnel potential to guide the dissociation pathway, combined with with umbrella sampling (details depicted in Figure S25), we calculated a binding free energy difference of (2.4 *±* 0.34) kcal/mol between the *wt*-primary state complex and the S189A mutant (Figure 5E). A similar calculation yielded a binding free energy difference of (1.2 *±* 0.43) kcal/mol between the *wt*-secondary state complex and the S189A mutant. These results indicate that the binding free energy difference between the *wt*-secondary state and the S189A mutant is approximately half that of the difference between *wt*-primary state and the S189A mutant, highlighting the greater conformational similarity of the secondary state to S189A compared to the primary state. Consistent with these findings, the binding free energy measured by fluorescence polarization (FP) falls between the two computed free energy bounds (Figure 5E), demonstrating strong agreement between simulation and experiment. Taken together, while our FP measurements and biased MD simulations demonstrate a reduced binding affinity upon the S189A mutation, NMR experiments reveal that this mutation does not affect MBD2’s selectivity for mCpG.

## 4 Discussion

In this study, we elucidated the atomic-level dynamics and stability of MBD2 binding to methylated CpG (mCpG) sites using extensive microsecond-scale MD simulations, validated by NMR spectroscopy and fluorescence polarization assays.

Initially, we assessed the stability of the MBD2–mCpG complex using its experimentally determined X-ray (PDB ID: 7MWK) and NMR (PDB ID: 2KY8^30^) structures. Using the X-ray structure as the initial conformation for MD simulations, we observe that the MBD2–mCpG complex remains fully stable, maintaining both its overall conformation and key base-specific interactions. This stability is mediated by the key MBD2 residues R166 and R188, which act as a clamp to lock MBD2 onto the mCpG site by interacting with the bases of opposing methylated cytosines, while Y178 reinforces the structure through a strong water-mediated hydrogen bond interaction with the base of methylcytosine mC ^−^_1_. These residues govern both the selectivity and affinity of MBD2 for mCpG^21^. Additionally, DNA methylation is a key determinant of the high stability and specificity of MBD2–mCpG interactions.^98^

In contrast to the recognition simulations (based on the X-ray structure) the MBD2–mCpG complex in the interrogation simulations (based on the NMR structure) appears unstable. This instability arises from the fact that the DNA in the NMR structure is partially modeled on canonical B-DNA due to low proton density observed during structure determination.^30^ Notably, despite the presence of key protein–DNA direct hydrogen bonds in the NMR structure, only three out of eight MD replicas (with a 1 *µ*s simulation time per replica) resemble the specific state observed in the X-ray structure, underscoring the significance of the indirect (shape) readout mechanism^99–101^ for stabilizing the specific complex. Figures 4A illustrate that a unique DNA conformation is necessary for the formation of the specific MBD2–mCpG complex. To adopt such unique DNA conformation required for proper recognition, MBD2—as shown in the NMR based interrogation simulations—must overcome an energetic barrier, which, based on the calculated similarity index^44^, is of moderate magnitude.

Consistent with these findings, our search simulations, for which MBD2 was initially positioned one base pair downstream of the mCpG site, reveal a moderate energetic barrier to recognition. In four out of seven replicas, MBD2 reaches the specific state observed in the X-ray structure on a timescale of several microseconds. Strikingly, even from a nonspecific B-DNA starting structure and with MBD2 positioned one base pair away from its specific target site, fully unbiased MD simulations spontaneously drive MBD2 to its native recognition complex, demonstrating the remarkable capability of MD simulations to recapitulate precise, functionally relevant protein–DNA recognition.

In three independent search simulations, MBD2 formed a stable alternative complex with mCpG, structurally distinct from the primary recognition state and remarkably persistent over multiple microseconds—2 up to about 35 *µ*s—highlighting its intrinsically stable nature. To uncover the structural determinants of the secondary state, we analyzed possible macro-switches distinguishing it from the primary binding mode. Our data indicate that the essential arginines R166 and R188 maintain their base-specific recognition of mCpG, suggesting a preserved core of selective interactions. Disruption of either residue has been experimentally shown to substantially impair MBD2’s affinity for methylated CpG^16,30^. In addition, the critical contact between K174 and a flanking mCpG base persists in both conformational states. Y178, which has also been associated with mCpG selectivity in MBD2, forms a water-mediated hydrogen bond in the primary state, whereas in the secondary state it interacts primarily through van der Waals interactions. Therefore, both states retain a core set of key interactions that robustly support MBD2’s binding selectivity and functional engagement with mCpG.

The key macro-switch differentiating the primary and secondary state is the interaction of S189 with the backbone of methylated cytosine mC_+1_. We observe that the absence of this interaction shifts the MBD2– mCpG conformation toward the secondary state. To investigate the precise role of S189 in mediating selective mCpG recognition by MBD2, we conducted ^15^N HSQC NMR spectroscopy on the MBD2–mCpG complex containing the S189A mutation and directly compared the resulting spectra with those of the wild-type complex.^77^ Importantly, our NMR analysis reveals that the S189A mutation does not compromise the selective recognition of mCpG by MBD2 (Figure 5C). However, both our fluorescence polarization experiments and biased MD simulations consistently show that the S189A mutation reduces MBD2’s binding affinity for mCpG (Figure 5D,E). Similar muations in MBD1 (S45A) and MeCP2 (S134A) have been shown to decrease their affinity for mCpG by approximately half.^31,32^ The S134C mutation in MeCP2 also reduces binding affinity for mCpG without affecting selectivity, and is even clinically significant, as it is associated with Rett syndrome, underscoring the biological importance of this serine residue in maintaining proper epigenetic regulation.^102^ We suggest that, analogous to S189A in MBD2, the S134C mutation could drive MeCP2 (whose conformation closely resembles that of MBD2, see Figure S26) into a binding conformation that diverges from its canonical state, potentially carrying greater functional consequences.

From our MD simulations of the MBD2–S189A variant, we observe that the S189A mutation indeed shifts the MBD2–mCpG complex toward a conformation characteristic of the secondary state. This finding strongly suggests that the secondary state represents a valid and selective recognition state, albeit with reduced binding affinity relative to the primary state.

To further establish the secondary state as a genuine alternative sequence-specific bound conformation, we conducted additional search simulations using a native DNA sequence. In line with previous results, bistability is recapitulated, with the secondary state closely resembling that observed with the initial DNA sequence.

Such bistable binding behavior in DNA-binding proteins were initially identified through single-molecule spectroscopy by Wenzhe Liu et al., demonstrating that the DNA-binding domain of the *Arabidopsis* WRKY protein undergoes bistable oscillations upon specific target engagement.^103^ Their findings further revealed that the dwell time, defined as the period the system resides in each oscillatory state, varies with temperature.^103^ Therefore, as X-ray crystallography preferentially captures well-ordered conformational states, the observation of only the primary state—being more ordered and less dynamic than the secondary state—may reflect this inherent crystallographic bias and may also explain why the complete structure could not be fully resolved in the NMR ensemble, owing to the dynamic nature of the MBD2–mCpG complex.^30,104,105^

It is also noteworthy that, since the S189–mC_+1_ interaction serves as a critical macro-switch distinguishing the two stable states observed with MBD2 bound at the target site, external factors could plausibly shift the equilibrium between these states by modulating the strength of this interaction. For instance, ions have been shown to modulate structural transitions in DNA-binding proteins^106^. Specifically, changes in ion concentrations regulate electrostatic interactions, particularly with the DNA backbone, which are critical for protein–DNA recognition^106–108^. Thus, variations in ionic strength may influence the S189–mC_+1_ backbone interaction, thereby increasing the likelihood of transitions between the two distinct stable states. Moreover, MBD2 binding to mCpG, in either the primary or secondary state, induces distinct DNA conformations. Our simulations reveal that disruption of the S189–mC_+1_ interaction leads to a distinct BI/BII backbone conformation at the mCpG site, which plays a critical role in determining the overall DNA geometry.^87–93^ Notably, MBD2 has been shown to retain high-affinity binding to mCpG across diverse DNA geometries—including both straight and bent conformations—suggesting that bistable recognition may balance conformational adaptability with high-affinity DNA binding. It has been shown that MBD2 becomes trapped in mCpG-rich genomic regions, yet can still rapidly exchange between closely spaced mCpG sites.^21^ In this context, the presence of an alternative bound state may provide a structural basis for maintaining specific mCpG recognition while enabling local conformational flexibility that facilitates exchange between neighboring methylated sites. Specifically, this bistable state mechanism may provide MBD2 with a strategic advantage, ensuring precise mCpG binding while also enabling dynamic participation in assembling larger complexes, such as MBD2–NuRD.

## 5 Conclusions

As a pivotal epigenetic regulator, MBD2 serves as a key reader of 5-methylcytosine (5mC), characterized by its high selectivity and tight binding to methylated CpG dinucleotides. To gain a deeper atomic-level understanding of the MBD2–mCpG complex, we performed extensive molecular dynamics simulations, both in its specific bound state and during transitions from nonspecific to specific binding. In summary, our study reveals the following key insights: (I) MBD2 adopts a bistable binding equilibrium with mCpG, a mechanism that may enable it to maintain high-affinity binding while remaining flexible for interactions across different molecular contexts and assemblies. (II) By favoring a conformation similar to the secondary state, the S189A mutation maintains MBD2’s selectivity for mCpG but compromises its binding affinity. (III) Our observation of bistable binding in MBD2 with two different DNA sequences, along with single-molecule evidence from the WRKY domain (Liu et al.), supports the notion that bound-state conformational equilibria may be a fundamental feature of sequence-specific protein–DNA recognition.

## Supporting information

MBD2_Manuscript_revision_Final_SI

## 6 Data Availability

The data that support the findings of this study are openly available at Zenodo (DOI: 10.5281/zenodo.15757877).

## 7 Acknowledgments

The authors thank the International Max Planck Research School for Biology and Computation (IMPRS-BAC), and the Bavarian Equal Opportunities Sponsorship—Realisierung von Chancengleichheit von Frauen in Forschung und Lehre (FFL)—for supporting S.V.. The authors gratefully acknowledge the computing time made available to them on the high-performance computer “Lise” at the NHR center NHR@ZIB. This center is jointly supported by the Federal Ministry of Education and Research and the state governments participating in the NHR (www.nhr-verein.de). The authors gratefully acknowledge the computing time provided by them on the high-performance computers Barnard and Romeo at NHR@TUD, which is funded by the Federal Ministry of Education and Research and the state governments participating on the basis of the resolutions of the GWK for the national high performance computing at universities (www.nhr-verein.de/unsere-partner). We thank Dustin Vivod for helpful discussions. This work was funded by the the Deutsche Forschungsgemeinschaft (DFG, German Research Foundation), project number 514664767, subproject B07 to M.E., project number IM141/1-3 to P.I., DFG RC Data Assimilation with Grant No. 318763901 to R.M., and National Institute of Diabetes, Digestive, and Kidney Diseases [R01DK115563 to D.W.].

## Author contributions

Senta Volkenandt (Conceptualization (lead), Data curation (lead), Formal analysis [lead], Investigation [lead], Methodology [lead], Software [lead], Validation [lead], Visualization [lead], Writing–original draft [supporting], Writing–review & editing [lead]), Julia Belyaeva (Data curation (supporting), Formal analysis [lead], Investigation [Supporrting], Methodology [lead], Software [lead], Validation [lead], Visualization [lead], Writing–original draft [supporting], Writing–review & editing [Supporting]), Torry Li (Formal analysis [Supporting], Methodology [Supporting], Validation [Supporting], Visualization [Supporting]), Matthias Elgeti (Visualization [Supporting], Writing–review & editing [Supporting], Supervision [Supporting]), Ralf Metzler (Conceptualization (Supporting), Supervision [Supporting], Writing–review & editing [Supporting]), Petra Imhof (Conceptualization (Supporting), Methodology [Supporting], Validation [Supporting], Resources [Supporting], Supervision [Supporting], Writing–review & editing [Supporting]), David C. Williams Jr. (Conceptualization (Supporting), Formal analysis [lead], Investigation [lead], Methodology [lead], Software [lead], Validation [lead], Visualization [lead], Resources [lead], Supervision [Supporting], Writing–original draft [supporting], Writing–review & editing [Supporting]), Mahdi Bagherpoor Helabad (Conceptualization (lead), Data curation (lead), Formal analysis [lead], Investigation [lead], Methodology [lead], Project Administration [lead], Resources [lead], Software [lead], Supervision [lead], Validation [lead], Visualization [lead], Writing–original draft [lead], Writing–review & editing [lead])

## References

[1] Luo, C., Hajkova, P., and Ecker, J. R. Dynamic DNA methylation: In the right place at the right time. Science. 2018; 361:1336–1340.

[2] Kanwal, R. and Gupta, S. Epigenetic modifications in cancer. Clin. Genet. 2012; 81:303–311.

[3] Moore, L. D., Le, T., and Fan, G. DNA methylation and its basic function. Neuropsychopharmacology. 2013; 38:23–38.

[4] Schübeler, D. Function and information content of DNA methylation. Nature. 2015; 517:321–326.

[5] Jiang, S. and Guo, Y. Epigenetic clock: DNA methylation in aging. Stem Cells Int. 2020; 2020:1047896.

[6] Smallwood, S. A. and Kelsey, G. De novo DNA methylation: a germ cell perspective. Trends Genet. 2012; 28:33–42.

[7] Smith, Z. D., Hetzel, S., and Meissner, A. DNA methylation in mammalian development and disease. Nat. Rev. Genet. 2024.

[8] Pasque, V., Jullien, J., Miyamoto, K., Halley-Stott, R. P., and Gurdon, J. Epigenetic factors influencing resistance to nuclear reprogramming. Trends Genet. 2011; 27:516–525.

[9] Blake, L. E., Roux, J., Hernando-Herraez, I., Banovich, N. E., Perez, R. G., Hsiao, C. J., Eres, I., Cuevas, C., Marques-Bonet, T., and Gilad, Y. A comparison of gene expression and DNA methylation patterns across tissues and species. Genome Res. 2020; 30:250–262.

[10] Hu, S., Wan, J., Su, Y., Song, Q., Zeng, Y., Nguyen, H. N., Shin, J., Cox, E., Rho, H. S., Woodard, C., et al. DNA methylation presents distinct binding sites for human transcription factors. eLife. 2013; 2:e00726.

[11] Bogdanović, O. and Veenstra, G. J. C. DNA methylation and methyl-CpG binding proteins: developmental requirements and function. Chromosoma. 2009; 118:549–565.

[12] Du, Q., Luu, P.-L., Stirzaker, C., and Clark, S. J. Methyl-CpG-binding domain proteins: readers of the epigenome. Epigenomics. 2015; 7:1051–1073.

[13] Hendrich, B. and Bird, A. Identification and characterization of a family of mammalian methyl CpG-binding proteins. Genet. Res. 1998; 72:59–72.

[14] Liu, K., Xu, C., Lei, M., Yang, A., Loppnau, P., Hughes, T. R., and Min, J. Structural basis for the ability of MBD domains to bind methyl-CG and TG sites in DNA. J. Biol. Chem. 2018; 293:7344–7354.

[15] Liu, K., Lei, M., Wu, Z., Gan, B., Cheng, H., Li, Y., and Min, J. Structural analyses reveal that MBD3 is a methylated CG binder. FEBS J. 2019; 286:3240–3254.

[16] Leighton, G. and Williams Jr, D. C. The methyl-CpG–Binding domain 2 and 3 proteins and formation of the nucleosome remodeling and deacetylase complex. J. Mol. Biol. 2020; 432:1624–1639.

[17] Cramer, J. M., Scarsdale, J. N., Walavalkar, N. M., Buchwald, W. A., Ginder, G. D., and Williams, D. C. Probing the dynamic distribution of bound states for methylcytosine-binding domains on DNA. J. Biol. Chem. 2014; 289:1294–1302.

[18] Zhang, Y., Ng, H.-H., Erdjument-Bromage, H., Tempst, P., Bird, A., and Reinberg, D. Analysis of the NuRD subunits reveals a histone deacetylase core complex and a connection with DNA methylation. Genes Dev. 1999; 13:1924–1935.

[19] Hainer, S. J., McCannell, K. N., Yu, J., Ee, L.-S., Zhu, L. J., Rando, O. J., and Fazzio, T. G. DNA methylation directs genomic localization of MBD2 and MBD3 in embryonic stem cells. eLife. 2016; 5:e21964.

[20] Wood, K. H. and Zhou, Z. Emerging molecular and biological functions of MBD2, a reader of DNA methylation. Front. Genet. 2016; 7:93.

[21] Leighton, G. O., Irvin, E. M., Kaur, P., Liu, M., You, C., Bhattaram, D., Piehler, J., Riehn, R., Wang, H., Pan, H., et al. Densely methylated DNA traps methyl-CpG–binding domain protein 2 but permits free diffusion by methyl-CpG–binding domain protein 3. J. Biol. Chem. 2022; 298.

[22] Mian, O. Y., Wang, S. Z., Zhu, S. Z., Gnanapragasam, M. N., Graham, L., Bear, H. D., and Ginder, G. D. Methyl-binding domain protein 2–dependent proliferation and survival of breast cancer cells. Mol. Cancer Res. 2011; 9:1152–1162.

[23] Lei, Q. and Zhang, W. Methyl CpG binding domain protein 2 (MBD2) in inflammation. Chin. Med. J. 2022; 135:2880–2882.

[24] Lax, E., Do Carmo, S., Enuka, Y., Sapozhnikov, D. M., Welikovitch, L. A., Mahmood, N., Rabbani, S. A., Wang, L., Britt, J. P., Hancock, W. W., et al. Methyl-CpG binding domain 2 (MBD2) is an epigenetic regulator of autism-risk genes and cognition. Transl. Psychiatry. 2023; 13:259.

[25] Baubec, T., Ivánek, R., Lienert, F., and Schübeler, D. Methylation-dependent and-independent genomic targeting principles of the MBD protein family. Cell. 2013; 153:480–492.

[26] Günther, K., Rust, M., Leers, J., Boettger, T., Scharfe, M., Jarek, M., Bartkuhn, M., and Renkawitz, R. Differential roles for MBD2 and MBD3 at methylated CpG islands, active promoters and binding to exon sequences. Nucleic Acids Res. 2013; 41:3010–3021.

[27] Yu, X., Azzo, A., Bilinovich, S. M., Li, X., Dozmorov, M., Kurita, R., Nakamura, Y., Williams Jr, D. C., and Ginder, G. D. Disruption of the MBD2-NuRD complex but not MBD3-NuRD induces high level HbF expression in human adult erythroid cells. Haematologica. 2019; 104:2361.

[28] Shang, S., Li, X., Azzo, A., Truong, T., Dozmorov, M., Lyons, C., Manna, A. K., Williams Jr, D. C., and Ginder, G. D. MBD2a-NuRD binds to the methylated-globin gene promoter and uniquely forms a complex required for silencing of HbF expression. Proc. Natl. Acad. Sci. U.S.A. 2023; 120:e2302254120.

[29] Millard, C. J., Fairall, L., Ragan, T. J., Savva, C. G., and Schwabe, J. W. The topology of chromatin-binding domains in the NuRD deacetylase complex. Nucleic Acids Res. 2020; 48:12972–12982.

[30] Scarsdale, J. N., Webb, H. D., Ginder, G. D., and Williams Jr, D. C. Solution structure and dynamic analysis of chicken MBD2 methyl binding domain bound to a target-methylated DNA sequence. Nucleic Acids Res. 2011; 39:6741–6752.

[31] Ohki, I., Shimotake, N., Fujita, N., Nakao, M., and Shirakawa, M. Solution structure of the methyl-CpG-binding domain of the methylation-dependent transcriptional repressor MBD1. EMBO J. 1999; 18:6653–6661.

[32] Free, A., Wakefield, R. I., Smith, B. O., Dryden, D. T., Barlow, P. N., and Bird, A. P. DNA recognition by the methyl-CpG binding domain of MeCP2. J. Biol. Chem. 2001; 276:3353–3360.

[33] Cheadle, J. P., Gill, H., Fleming, N., Maynard, J., Kerr, A., Leonard, H., Krawczak, M., Cooper, D. N., Lynch, S., Thomas, N., et al. Long-read sequence analysis of the MeCP2 gene in Rett syndrome patients: correlation of disease severity with mutation type and location. Hum. Mol. Genet. 2000; 9:1119–1129.

[34] Kucukkal, T. G., Yang, Y., Uvarov, O., Cao, W., and Alexov, E. Impact of Rett syndrome mutations on MeCP2 MBD stability. Biochemistry. 2015; 54:6357–6368.

[35] Walavalkar, N. M., Cramer, J. M., Buchwald, W. A., Scarsdale, J. N., and Williams Jr, D. C. Solution structure and intramolecular exchange of methyl-cytosine binding domain protein 4 (MBD4) on DNA suggests a mechanism to scan for mCpG/TpG mismatches. Nucleic Acids Res. 2014; 42:11218–11232.

[36] Pan, H., Bilinovich, S. M., Kaur, P., Riehn, R., Wang, H., and Williams Jr, D. C. CpG and methylation-dependent DNA binding and dynamics of the methylcytosine binding domain 2 protein at the single-molecule level. Nucleic Acids Res. 2017; 45:9164–9177.

[37] Bigman, L. S., Greenblatt, H. M., and Levy, Y. What are the molecular requirements for protein sliding along DNA?. J. Phys. Chem. B. 2021; 125:3119–3131.

[38] Blainey, P. C., Luo, G., Kou, S., Mangel, W. F., Verdine, G. L., Bagchi, B., and Xie, X. S. Nonspecifically bound proteins spin while diffusing along DNA. Nat. Struct. Mol. Biol. 2009; 16:1224–1229.

[39] Marklund, E., van Oosten, B., Mao, G., Amselem, E., Kipper, K., Sabantsev, A., Emmerich, A., Globisch, D., Zheng, X., Lehmann, L. C., et al. DNA surface exploration and operator bypassing during target search. Nature. 2020; 583:858–861.

[40] Cencini, M. and Pigolotti, S. Energetic funnel facilitates facilitated diffusion. Nucleic Acids Res. 2018; 46:558–567.

[41] Tempestini, A., Monico, C., Gardini, L., Vanzi, F., Pavone, F. S., and Capitanio, M. Sliding of a single lac repressor protein along DNA is tuned by DNA sequence and molecular switching. Nucleic Acids Res. 2018; 46:5001–5011.

[42] Ahmadi, A., Rosnes, I., Blicher, P., Diekmann, R., Schüttpelz, M., Glette, K., Tørresen, J., Bjørs, M., Dalhus, B., and Rowe, A. D. Breaking the speed limit with multimode fast scanning of DNA by Endonuclease V. Nat. Commun. 2018; 9:5381.

[43] Arbel-Goren, R., McKeithen-Mead, S. A., Voglmaier, D., Afremov, I., Teza, G., Grossman, A. D., and Stavans, J. Target search by an imported conjugative DNA element for a unique integration site along a bacterial chromosome during horizontal gene transfer. Nucleic Acids Res. 2023; 51:3116–3129.

[44] Leven, I. and Levy, Y. Quantifying the two-state facilitated diffusion model of protein-DNA interactions. Nucleic Acids Res. 2019; 47:5530–5538.

[45] Marcovitz, A. and Levy, Y. Frustration in protein-DNA binding influences conformational switching and target search kinetics. Proc. Natl. Acad. Sci. U.S.A. 2011; 108:17957–17962.

[46] Pulkkinen, O. and Metzler, R. Distance matters: the impact of gene proximity in bacterial gene regulation. Phys. Rev. Lett. 2013; 110:198101.

[47] Hedström, L., Metzler, R., Lizana, L. Enhancer-insulator pairing reveals heterogeneous dynamics in long-distance 3D gene regulation. PRX Life. 2024;2(3):033008.

[48] Marklund, E. G., Mahmutovic, A., Berg, O. G., Hammar, P., van der Spoel, D., Fange, D., and Elf, J. Transcription-factor binding and sliding on DNA studied using micro-and macroscopic models. Proc. Natl. Acad. Sci. U.S.A. 2013; 110:19796–19801.

[49] Terakawa, T. and Takada, S. p53 dynamics upon response element recognition explored by molecular simulations. Sci. Rep. 2015; 5:1–10.

[50] Bhattacherjee, A., Krepel, D., and Levy, Y. Coarse-grained models for studying protein diffusion along DNA. Wiley Interdiscip. Rev. Comput. Mol. Sci. 2016; 6:515–531.

[51] Chu, X. and Munoz, V. Roles of conformational disorder and downhill folding in modulating protein–DNA recognition. Phys. Chem. Chem. Phys. 2017; 19:28527–28539.

[52] Wieczór, M. and Czub, J. How proteins bind to DNA: target discrimination and dynamic sequence search by the telomeric protein TRF1. Nucleic Acids Res. 2017; 45:7643–7654.

[53] Zacharias, M. Atomic resolution insight into Sac7d protein binding to DNA and associated global changes by molecular dynamics simulations. Angew. Chem. Int. Ed. 2019; 58:5967–5972.

[54] Tian, J., Wang, L., and Da, L.-T. Atomic resolution of short-range sliding dynamics of thymine DNA glycosylase along DNA minor-groove for lesion recognition. Nucleic Acids Res. 2021; 49:1278–1293.

[55] Dai, L., Xu, Y., Du, Z., Su, X.-D., and Yu, J. Revealing atomic-scale molecular diffusion of a plant-transcription factor WRKY domain protein along DNA. Proc. Natl. Acad. Sci. U.S.A. 2021; 118:e2102621118.

[56] Pettersen, E. F., Goddard, T. D., Huang, C. C., Couch, G. S., Greenblatt, D. M., Meng, E. C., and Ferrin, T. E. UCSF Chimera—a visualization system for exploratory research and analysis. J. Comput. Chem. 2004; 25:1605–1612.

[57] Abraham, M. J., Murtola, T., Schulz, R., Páll, S., Smith, J. C., Hess, B., and Lindahl, E. GROMACS: High performance molecular simulations through multi-level parallelism from laptops to supercomputers. SoftwareX. 2015; 1:19–25.

[58] Maier, J. A., Martinez, C., Kasavajhala, K., Wickstrom, L., Hauser, K. E., and Simmerling, C. ff14SB: improving the accuracy of protein side chain and backbone parameters from ff99SB. Journal of Chemical Theory and Computation. 2015; 11:3696–3713.

[59] Ivani, I., Dans, P. D., Noy, A., Pérez, A., Faustino, I., Hospital, A., Walther, J., Andrio, P., Goni, R., Balaceanu, A., et al. Parmbsc1: a refined force field for DNA simulations. Nat. Methods. 2016; 13:55–58.

[60] Liebl, K. and Zacharias, M. How methyl-sugar interactions determine DNA structure and flexibility. Nucleic Acids Res. 2019; 47:1132–1140.

[61] Carvalho, A. T., Gouveia, L., Kanna, C. R., Wärmländer, S. K., Platts, J. A., and Kamerlin, S. C. L. Understanding the structural and dynamic consequences of DNA epigenetic modifications: Computational insights into cytosine methylation and hydroxymethylation. Epigenetics. 2014; 9:1604–1612.

[62] Jorgensen, W. L., Chandrasekhar, J., Madura, J. D., Impey, R. W., and Klein, M. L. Comparison of simple potential functions for simulating liquid water. J. Chem. Phys. 1983; 79:926–935.

[63] Bussi, G., Donadio, D., and Parrinello, M. Canonical sampling through velocity rescaling. J. Chem. Phys. 2007; 126:014101.

[64] Parrinello, M. and Rahman, A. Polymorphic transitions in single crystals: A new molecular dynamics method. J. Appl. Phys. 1981; 52:7182–7190.

[65] Darden, T., York, D., and Pedersen, L. Particle mesh Ewald: An N · log(N) method for Ewald sums in large systems. J. Chem. Phys. 1993; 98:10089–10092.

[66] Essmann, U., Perera, L., Berkowitz, M. L., Darden, T., Lee, H., and Pedersen, L. G. A smooth particle mesh Ewald method. J. Chem. Phys. 1995; 103:8577–8593.

[67] Hess, B., Bekker, H., Berendsen, H. J., and Fraaije, J. G. LINCS: a linear constraint solver for molecular simulations. J. Comput. Chem. 1997; 18:1463–1472.

[68] PLUMED consortium. Promoting transparency and reproducibility in enhanced molecular simulations. Nat. Methods. 2019; 16:670–673.

[69] Limongelli, V., Bonomi, M., and Parrinello, M. Funnel metadynamics as accurate binding free-energy method. Proc. Natl. Acad. Sci. U.S.A. 2013; 110:6358–6363.

[70] Schrodinger, LLC. The PyMOL molecular graphics system. Version. 2015; 1:8.

[71] Grubmüller, H., Heymann, B., and Tavan, P. Ligand binding: molecular mechanics calculation of the streptavidin-biotin rupture force. Science. 1996; 271:997–999.

[72] Van Der Spoel, D., Lindahl, E., Hess, B., Groenhof, G., Mark, A. E., and Berendsen, H. J. GROMACS: fast, flexible, and free. J. Comput. Chem. 2005; 26:1701–1718.

[73] Tribello, G. A., Bonomi, M., Branduardi, D., Camilloni, C., and Bussi, G. PLUMED 2: New feathers for an old bird. Comput. Phys. Commun. 2014; 185:604–613.

[74] Torrie, G. M. and Valleau, J. P. Nonphysical sampling distributions in Monte Carlo free-energy estimation: Umbrella sampling. J. Comput. Phys. 1977; 23:187–199.

[75] Kumar, S., Rosenberg, J. M., Bouzida, D., Swendsen, R. H., and Kollman, P. A. The weighted histogram analysis method for free-energy calculations on biomolecules. I. The method. J. Comput. Chem. 1992; 13:1011–1021.

[76] Souaille, M. and Roux, B. Extension to the weighted histogram analysis method: combining umbrella sampling with free energy calculations. Comput. Phys. Commun. 2001; 135:40–57.

[77] Desai, M. A., Webb, H. D., Sinanan, L. M., Scarsdale, J. N., Walavalkar, N. M., Ginder, G. D., and Williams Jr, D. C. An intrinsically disordered region of methyl-CpG binding domain protein 2 (MBD2) recruits the histone deacetylase core of the NuRD complex. Nucleic Acids Res. 2015; 43:3100–3113.

[78] Cai, M., Williams, D. C., Wang, G., Lee, B. R., Peterkofsky, A., and Clore, G. M. Solution structure of the phosphoryl transfer complex between the signal-transducing protein IIAGlucose and the cytoplasmic domain of the glucose transporter IICBGlucose of the Escherichia coli glucose phosphotransferase system. J. Biol. Chem. 2003; 278:25191–25206.

[79] Delaglio, F., Grzesiek, S., Vuister, G. W., Zhu, G., Pfeifer, J., and Bax, A. NMRPipe: a multidimensional spectral processing system based on UNIX pipes. J. Biomol. NMR. 1995; 6:277–293.

[80] Vranken, W. F., Boucher, W., Stevens, T. J., Fogh, R. H., Pajon, A., Llinas, M., Ulrich, E. L., Markley, J. L., Ionides, J., and Laue, E. D. The CCPN data model for NMR spectroscopy: development of a software pipeline. Proteins. 2005; 59:687–696.

[81] Best, R. B., Hummer, G., and Eaton, W. A. Native contacts determine protein folding mechanisms in atomistic simulations. Proc. Natl. Acad. Sci. U.S.A. 2013; 110:17874–17879.

[82] Daura, X., Gademann, K., Jaun, B., Seebach, D., van Gunsteren, W. F., and Mark, A. E. Peptide Folding: When Simulation Meets Experiment. Angew. Chem. Int. Ed. 1999; 38:236–240.

[83] Hess, B. Convergence of sampling in protein simulations. Phys. Rev. E. 2002; 65:031910.

[84] Lavery, R., Moakher, M., Maddocks, J. H., Petkeviciute, D., and Zakrzewska, K. Conformational analysis of nucleic acids revisited: Curves+. Nucleic Acids Res. 2009; 37:5917–5929.

[85] Blanchet, C., Pasi, M., Zakrzewska, K., and Lavery, R. CURVES+ web server for analyzing and visualizing the helical, backbone and groove parameters of nucleic acid structures. Nucleic Acids Res. 2011; 39:W68–W73.

[86] Eisenhaber, F., Lijnzaad, P., Argos, P., Sander, C., and Scharf, M. The double cubic lattice method: Efficient approaches to numerical integration of surface area and volume and to dot surface contouring of molecular assemblies. J. Comput. Chem. 1995; 16:273–284.

[87] Djuranovic, D. and Hartmann, B. DNA fine structure and dynamics in crystals and in solution: the impact of BI/BII backbone conformations. Biopolymers. 2004; 73:356–368.

[88] Heddi, B., Foloppe, N., Bouchemal, N., Hantz, E., and Hartmann, B. Quantification of DNA BI/BII backbone states in solution. Implications for DNA overall structure and recognition. J. Am. Chem. Soc. 2006; 128:9170–9177.

[89] Tian, Y., Kayatta, M., Shultis, K., Gonzalez, A., Mueller, L. J., and Hatcher, M. E. 31P NMR investigation of backbone dynamics in DNA binding sites. J. Phys. Chem. B. 2009; 113:2596–2603.

[90] Westwood, M., Ljunggren, K., Boyd, B., Becker, J., Dwyer, T. J., and Meints, G. A. Single-base lesions and mismatches alter the backbone conformational dynamics in DNA. Biochemistry. 2021; 60:873–885.

[91] Westwood, M., Pilarski, A., Johnson, C., Mamoud, S., and Meints, G. Backbone conformational equilibrium in mismatched DNA correlates with enzyme activity. Biochemistry. 2023; 62:2816–2827.

[92] Savelyev, A. and MacKerell Jr, A. D. Differential deformability of the DNA minor groove and altered BI/BII backbone conformational equilibrium by the monovalent ions Li+, Na+, K+, and Rb+ via water-mediated hydrogen bonding. Journal of chemical theory and computation. 2015; 11:4473–4485.

[93] Precechtelová, J., Novák, P., Munzarová, M. L., Kaupp, M., and Sklenar, V. Phosphorus chemical shifts in a nucleic acid backbone from combined molecular dynamics and density functional calculations. J. Am. Chem. Soc. 2010; 132:17139–17148.

[94] Cramer, J. M., Cramer, J. M., Pohlmann, D., Pohlmann, D., Gómez, F., Mark, L., Kornegay, B., Hall, C., Siraliev-Perez, E., Walavalkar, N. M., Walavalkar, N. M., Sperlazza, M., Bilinovich, S. M., Prokop, J., Hill, A., and Williams, D. C. Methylation specific targeting of a chromatin remodeling complex from sponges to humans. Sci. Rep. 2017; 7:40674.

[95] Sperlazza, M. J., Bilinovich, S. M., Sinanan, L. M., Javier, F. R., and Williams, Jr, D. C. Structural Basis of MeCP2 Distribution on Non-CpG Methylated and Hydroxymethylated DNA. J. Mol. Biol. 2017; 429:1581–1594.

[96] Lee, H. R., Helquist, S. A., Kool, E. T., and Johnson, K. A. Importance of hydrogen bonding for efficiency and specificity of the human mitochondrial DNA polymerase. J. Biol. Chem. 2008; 283:14402–14410.

[97] Gorfe, A. A. and Jelesarov, I. Energetics of Sequence-Specific Protein-DNA Association: Computational Analysis of Integrase Tn916 Binding to Its Target DNA. Biochemistry. 2003; 42:11568–11576.

[98] Ginder, G. D. and Williams Jr, D. C. Readers of DNA methylation, the MBD family as potential therapeutic targets. Pharmacol. Ther. 2018; 184:98–111.

[99] Rohs, R., Jin, X., West, S. M., Joshi, R., Honig, B., and Mann, R. S. Origins of specificity in protein-DNA recognition. Annu. Rev. Biochem. 2010; 79:233–269.

[100] Samee, M. A. H., Bruneau, B. G., and Pollard, K. S. A de novo shape motif discovery algorithm reveals preferences of transcription factors for DNA shape beyond sequence motifs. Cell Syst. 2019; 8:27–42.

[101] Sangeeta Mishra, S. K., and Bhattacherjee, A. Role of Shape Deformation of DNA-Binding Sites in Regulating the Efficiency and Specificity in Their Recognition by DNA-Binding Proteins. JACS Au. 2024; 4:2640–2655.

[102] Yang, Y., Kucukkal, T. G., Li, J., Alexov, E., and Cao, W. Binding analysis of methyl-CpG binding domain of MeCP2 and Rett syndrome mutations. ACS Chem. Biol. 2016; 11:2706–2715.

[103] Liu, W., Li, J., Xu, Y., Yin, D., Zhu, X., Fu, H., Su, X., and Guo, X. Complete Mapping of DNA-Protein Interactions at the Single-Molecule Level. Adv. Sci. 2021; 8:2101383.

[104] Tilton Jr, R. F., Dewan, J. C., and Petsko, G. A. Effects of temperature on protein structure and dynamics: X-ray crystallographic studies of the protein ribonuclease-A at nine different temperatures from 98 to 320K. Biochemistry. 1992; 31:2469–2481.

[105] Fraser, J. S., Van Den Bedem, H., Samelson, A. J., Lang, P. T., Holton, J. M., Echols, N., and Alber, T. Accessing protein conformational ensembles using room-temperature X-ray crystallography. Proc. Natl. Acad. Sci. U.S.A. 2011; 108:16247–16252.

[106] Xu, C., Lu, Y., Wu, Y., Yuan, S., Ma, J., Fu, H., Wang, H., Wang, L., Zhang, H., Yu, X., et al. Sodium Ion-Induced Structural Transition on the Surface of a DNA-Interacting Protein. Adv. Sci. 2024; 11:2401838.

[107] Pasi, M., Maddocks, J. H., and Lavery, R. Analyzing ion distributions around DNA: sequence-dependence of potassium ion distributions from microsecond molecular dynamics. Nucleic Acids Res. 2015; 43:2412–2423.

[108] Yu, B., Pettitt, B. M., and Iwahara, J. Dynamics of ionic interactions at protein-nucleic acid interfaces. Acc. Chem. Res. 2020; 53:1802–1810.

