## Supplementary material for "Exploring the bistable equilibrium of methylated CpG DNA recognition by the MBD2 protein": MBD2_Manuscript_revision_Final_SI: MBD2_Manuscript_revision_Final_clean_SI.pdf

#### Additional information on supplementary figures

**Figure S1:** Recognition simulations are highly stable across all replicas, yielding an average RMSD of $1.84 \pm 0.11$ Å over the runs of the full trajectories and $1.87 \pm 0.12$ Å when considering only the 300–1000 ns interval (Figure S1**A**). In contrast, the interrogation simulations initiated from the NMR structure approach the equilibrated crystal conformation only in replicas I2 and I7, and toward the end of the trajectory in replica I3 (Figure S1**B**). The shaded area in Figure S1**B** shows two stardard deviations of the recognition simulations' fluctuations, meaning that $\sim 95\,\%$ of recognition simulation data after 300 ns can be found between 1.17 Å and 2.57 Å. Figure S1**C** shows that only replicas I1, I2, I3 and I7 have an average RMSD close to 3 Å, indicating that the interrogation simulations do not stay close to their initial structure in contrast to the recognition simulations. The higher structural flexibility of the interrogation simulations is reflected in a higher average average RMSD of $3.64 \pm 0.85$ Å, experiencing also a higher standard deviation.

**Figure S4 , S5**: Additional characterization of the search simulations and the initial 1D sliding of MBD2 around DNA.
The MBD2 protein is placed one base pair downstream from its target site (Figure S4). We can see that the system adopts a structure in agreement with recognition in four (S1, S2, S6, S7) of the seven replicas.
For the first measure the central eight base pairs of the reference model's DNA are aligned to the $x$–axis with their center of mass (COM) being placed in the origin at $(0, 0, 0)$. A trajectory can now be aligned to that DNA and the angle $\theta_1(t)$ in the $y$–$z$–plane of the COM of the protein's backbone can be calculated relative to the reference (Figure S4**B**).
For the second measure we calculated the angle $\theta_2(t)$ between the vector that is formed by the P-atoms

(COM) of the opposing mCpG dinucleotides and the beta sheet. In more detail, the first vector is defined between the COM of the P atoms of $mC_{-1}$ and $G_{+1}$ in one DNA strand and the COM of the P atoms of $G_{-1}$ and $mC_{+1}$ in the other DNA strand. The second vector is defined between the COM of the C$\alpha$ atoms of V164 and D176, and the COM of the C$\alpha$ atoms of K160 and F180 (Figure S4**C**, C$\alpha$ and P atoms are shown as black spheres).
From Figure S5 we can see that the initial sliding happens on the sub-microsecond timescale. The beta-sheet can be in the correct orientation relative to the mCpG dinucleotide for the first time even within the first few nanoseconds but latest after $\sim 40$ ns. For MBD2's COM it can take longer to approach the desired orientation for the first time (e.g. S1) or it can be on the same timescale (e.g. S2). Importantly, the protein can only stay stably at the recognition site if corresponding interactions are formed. We notice that the three replicas that do not end up in the recognition state, still exhibit stable orientation behaviour. There, $\theta_1$ is about $9-13°$ lower and $\theta_2$ $7-12°$ higher, which can be explained by the different hydrogen bond pattern of the two WT clusters.

**Figure S6 , S7 , S11:** RMSD clustering and validation
The distribution of pairwise RMSD values was fitted with a sum of scaled normal distribution functions of the form $\frac{A}{\sqrt{2\pi\sigma^2}} \exp\left(-\frac{(x-\mu)^2}{2\sigma^2}\right)$, with parameters $A$, $\mu$, and $\sigma$. Three components were used for the wild-type (WT) simulations (Figure S6**A**) and two components for the S189A mutant simulations (Figure S6**B**). The WT dataset includes all recognition, interrogation, and search simulations.
For the WT simulations, the RMSD distribution supports meaningful clustering, as the fitted distribution shows a clear valley separating lower- and higher-RMSD populations. The first fitted peak, centered at $1.85$ Å, coincides with the average RMSD of the recognition simulations. We therefore selected a natural clustering threshold of $\mu + 2\sigma \approx 2.5$ Å.
This choice is further supported by principal component analysis (PCA). The first two principal components (PCs) capture $\sim 60\%$ of the variance, while the first three PCs capture approximately two-thirds of the variance. Their cosine contents are $0.002$, $0.002$, and $0.008$, respectively, indicating that these PCs are not dominated by random diffusion. The transition between the primary and secondary states is clearly captured along PC1 (Figure S7), where cluster 2 (Figure S7**B**) is broader than cluster 1 (Figure S7**A**).
Because sequence-specific DNA recognition is defined by the MBD2–DNA interface, we also analyzed the contact area between MBD2 and DNA. The absence of multimodal distributions in the WT contact-area profiles (Figure S11**A**) further supports the selected RMSD threshold.
In contrast to the WT simulations, the RMSD distribution of the S189A mutant simulations shows only a shoulder rather than a clear separation (Figure S6**B**), suggesting that the sampled conformations do not form well-separated states. Thus, the resulting clusters depend more strongly on the chosen threshold and should be interpreted as indicators of trajectory evolution rather than distinct conformational states. We selected a threshold of $2.2$ Å because it yields approximately normal contact-area distributions (Figure S11**B**). In contrast, higher thresholds lead to bimodal distributions (data not shown), making the clustering less suitable for analyzing protein–DNA interactions.

### Supplementary tables

**Table S1.** System setups in MD simulations (Force Field: AMBER14SB_BSC1_TIP3P)

| System | $x \times y \times z$ [1] | $N_{\mathrm{atoms}}$ [2] | $N_{\mathrm{H_2O}}$ [3] | $N_{\mathrm{ion}}$ [4] | $N_{\mathrm{rep}}$ [5] | $\tau_s$ [6] |
|---|---|---|---|---|---|---|
| Search | 8.30×8.30×8.30 | 57 124 | 18 271 | 59 | 7 | 134 $\mu$s |
| Search DNA2 | 8.30×8.30×8.30 | 56 954 | 18 219 | 59 | 9 | 90 $\mu$s |
| Recognition | 8.30×8.30×8.30 | 56 904 | 18 257 | 59 | 8 | 8 $\mu$s |
| Interrogation | 8.26×8.26×8.26 | 57 026 | 18 282 | 55 | 8 | 8 $\mu$s |
| S189A | 8.14×8.14×8.14 | 52 844 | 16 966 | 53 | 8 | 24 $\mu$s |
| Steered MD (primary state) | 8.30×8.30×8.30 | 56 904 | 18 257 | 59 | 20 | 1.6 $\mu$s |
| Steered MD (secondary state) | 8.30×8.30×8.30 | 57 124 | 18 271 | 59 | 20 | 1.6 $\mu$s |
| Steered MD (S189A) | 8.14×8.14×8.14 | 52 844 | 16 966 | 53 | 20 | 1.6 $\mu$s |
| Umbrella sampling (primary state) | 8.30×8.30×8.30 | 56 904 | 18 257 | 59 | 35 | 2.8 $\mu$s |
| Umbrella sampling (secondary state) | 8.30×8.30×8.30 | 57 124 | 18 271 | 59 | 35 | 2.8 $\mu$s |
| Umbrella sampling (S189A) | 8.14×8.14×8.14 | 52 844 | 16 966 | 53 | 35 | 2.8 $\mu$s |
| Total | | | | | | 277.2 $\mu$s |

**Table S2.** Pairwise RMSD values (in Å) are shown for the MBD2 crystal structure (PDB ID: 7MWK), the reference model (equilibrated crystal structure), and the medoid structure of the recognition simulations (R). The medoid structure is defined as the structure closest to the ensemble average based on the C$\alpha$ atoms of the protein core and the P atoms of DNA. Both the reference model and the medoid structure closely resemble the X-ray structure.

| | Crystal | Equil. crystal | Medoid (R) |
|---|---|---|---|
| Crystal | 0 | 1.04 | 1.65 |
| Equil. crystal | 1.04 | 0 | 1.35 |
| Medoid (R) | 1.65 | 1.35 | 0 |

[1] Initial simulation box size ($nm^3$).
[2] Total number of atoms.
[3] Total number of water molecules.
[4] Total number of (counter / salt) ions.
[5] Total number of replicas.
[6] Total simulations time.

**Table S3.** RMSD values (in Å) between RMSD cluster centers. Best fits between the reference model (equilibrated crystal structure) and the wild-type (WT) and S189A clusters are highlighted in bold. WT cluster $C_1^{wt}$, S189A cluster $C_1^{mu}$, and the reference model, are structurally very similar. S189A clusters $C_2^{mu}$ and $C_3^{mu}$ are closest to WT cluster $C_2^{wt}$.

| | | reference model | WT | | S189A | | |
|---|---|---|---|---|---|---|---|
| | | | $C_1^{wt}$ | $C_2^{wt}$ | $C_1^{mu}$ | $C_2^{mu}$ | $C_3^{mu}$ |
| Equil. cryst. | | 0 | **1.8** | 3.6 | **1.4** | 3.0 | 3.6 |
| WT | $C_1^{wt}$ | **1.8** | 0 | 4.1 | **1.1** | 3.2 | 4.2 |
| | $C_2^{wt}$ | 3.6 | 4.1 | 0 | 3.7 | **2.2** | **2.5** |
| S189A | $C_1^{mu}$ | **1.4** | **1.1** | 3.7 | 0 | 2.9 | 3.8 |
| | $C_2^{mu}$ | 3.0 | 3.2 | **2.2** | 2.9 | 0 | 3.1 |
| | $C_3^{mu}$ | 3.6 | 4.2 | **2.5** | 3.8 | 3.1 | 0 |

**Table S4.** RMSD values in Å between the RMSD cluster centers of the simulations wild-type MBD2 simulations with different DNA sequences DNA 1 and DNA 2 as shown in Figure 4C. Best fits between the two clustering are highlighted in bold.

| | | WT DNA 1 | |
|---|---|---|---|
| | | $C_1^{wt}$ | $C_2^{wt}$ |
| WT DNA 2 | $C_1^{wt,\mathrm{N}}$ | **2.6** | 5.0 |
| | $C_2^{wt,\mathrm{N}}$ | 3.9 | **1.4** |

#### Supplementary figures

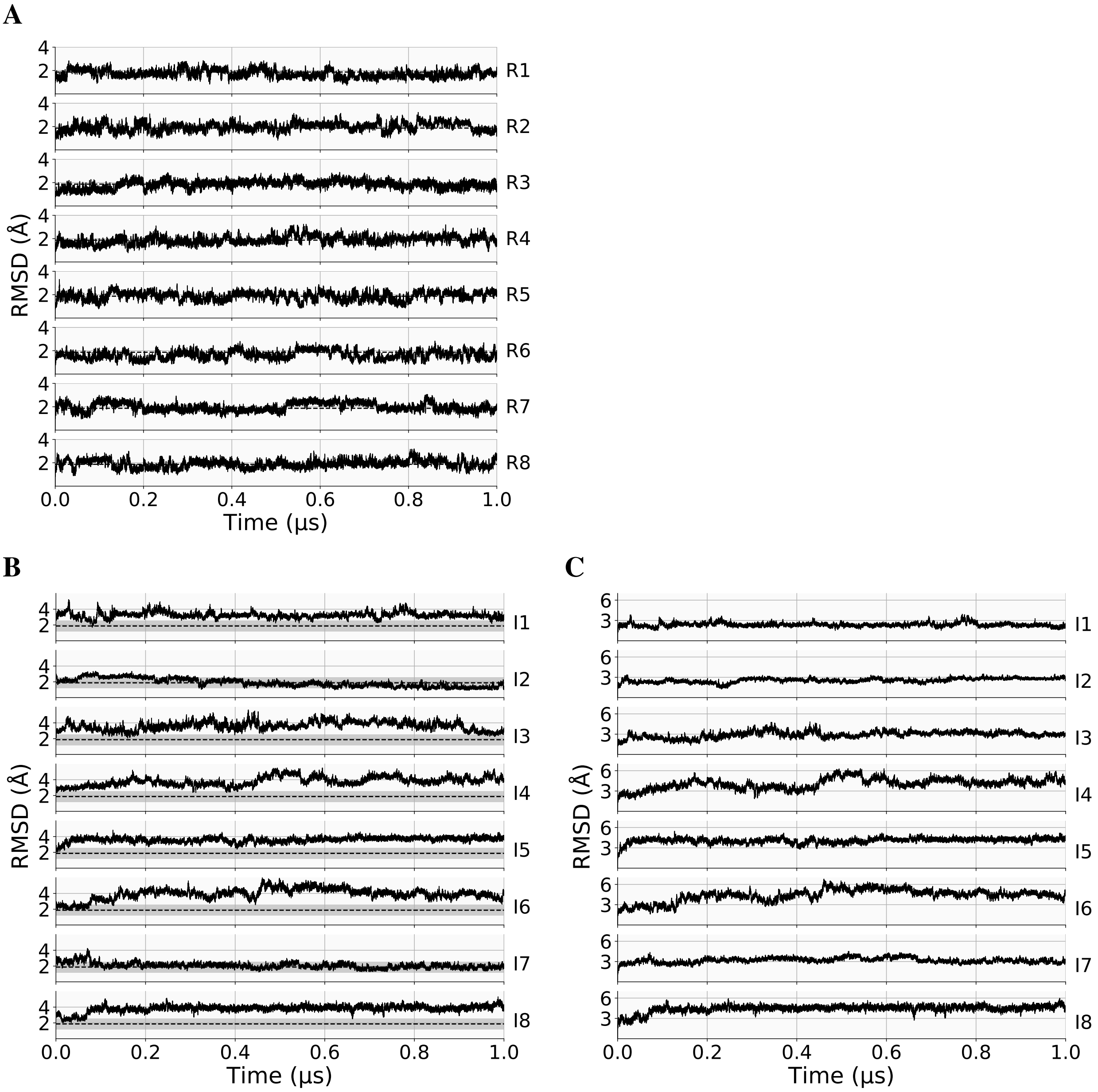


**Figure S1.** RMSD of the **A** recognition (R) and **B** interrogation (I) simulations with respect to the reference model based on the X-ray structure. The average RMSD $1.87$ Å $(300 - 1000\,\mathrm{ns})$ of the recognition simulations is highlighted as a dashed line. The shaded area in **B** additionally indicates the fluctuation around that average as two standard deviations. **C** RMSD of the interrogation simulations with respect to the first frame. Note the different RMSD scales

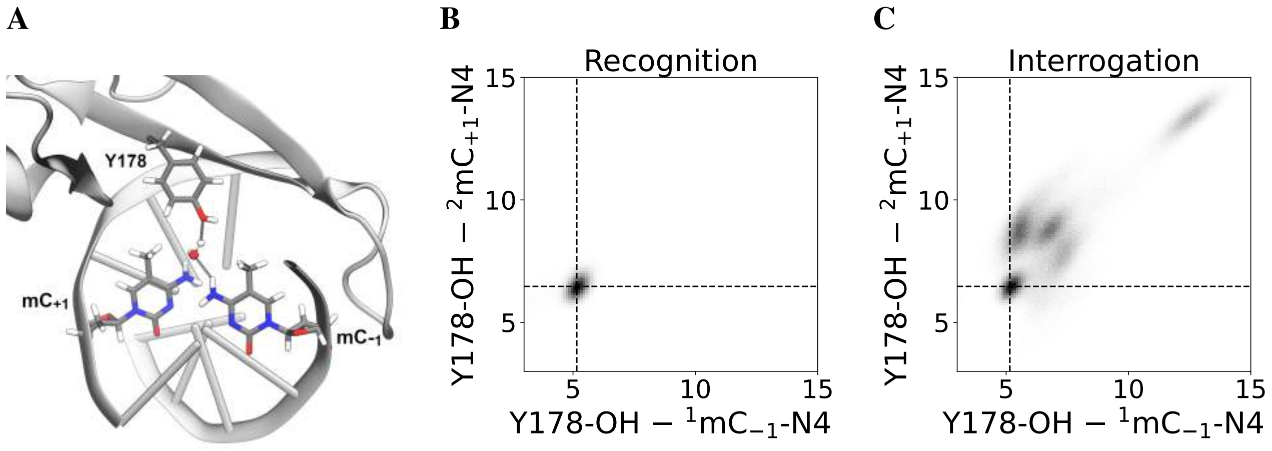


**Figure S2.** **A** Water-mediated hydrogen bond of Y178 with $mC_{-1}$ in the reference model. Distribution of the distances between the OH-atom of Y178 and the N4-atom of the methylated cytosines for the **B** recognition and **C** interrogation simulations. Dashed lines indicate the average values of the recognition simulations.

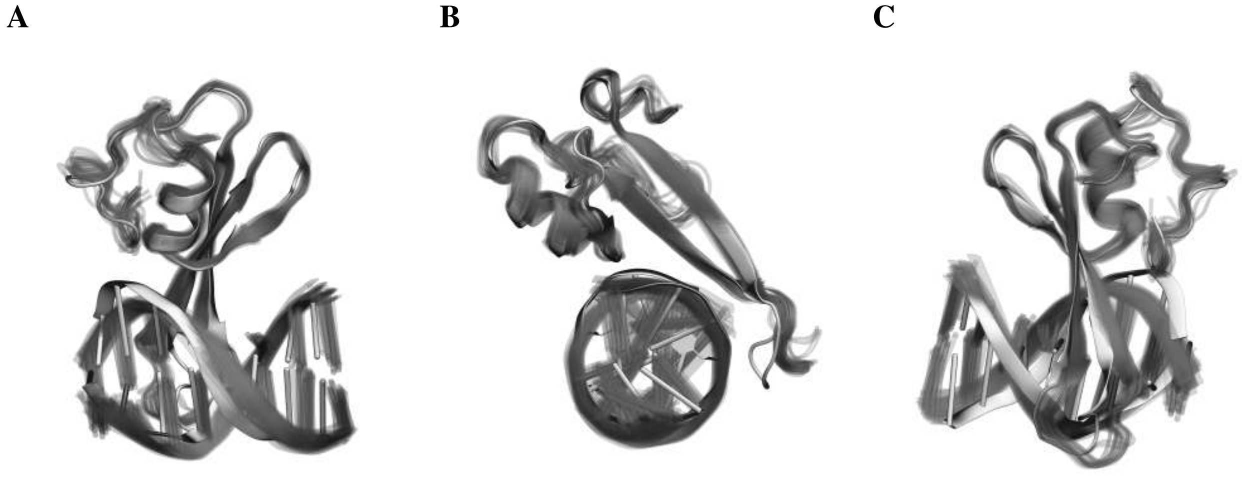


**Figure S3.** Overlay of the core region of crystal (PDB ID: 7MWK) and NMR (PDB ID: 2KY8) structure of MBD2 in solid and transparent, respectively, in **A** front, **B** side, and **C** back view. The DNA in the NMR structures adopts a different conformation, as it was modeled as B-DNA. This leads to a larger separation between the DNA and the MBD2 head domain, including helix $\alpha 1$. The tail domain is also displaced further from the DNA, as most clearly seen in **B**.

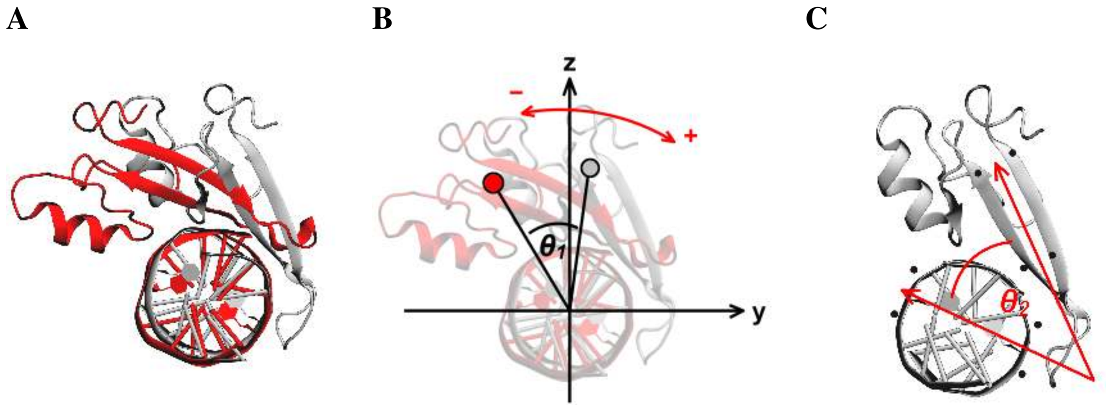


**Figure S4.** **A** Initial structure of the search simulations (red) aligned to the DNA of the reference model (grey). **B** Visualisations of the angle $\theta_1(t)$ of the protein backbone COM around the DNA, which is aligned to the $x$–axis. **C** Visualisation of $\theta_2$ and the two vectors defined by the mCpG dinucleotide and the beta-sheet; atoms used for COMs are highlighted as black spheres. In more detail, the first vector is defined between the COM of the P atoms of $mC_{-1}$ and $G_{+1}$ in one DNA strand and the COM of the P atoms of $G_{-1}$ and $mC_{+1}$ in the other DNA strand. The second vector is defined between the COM of the C$\alpha$ atoms of V164 and D176, and the COM of the C$\alpha$ atoms of K160 and F180.

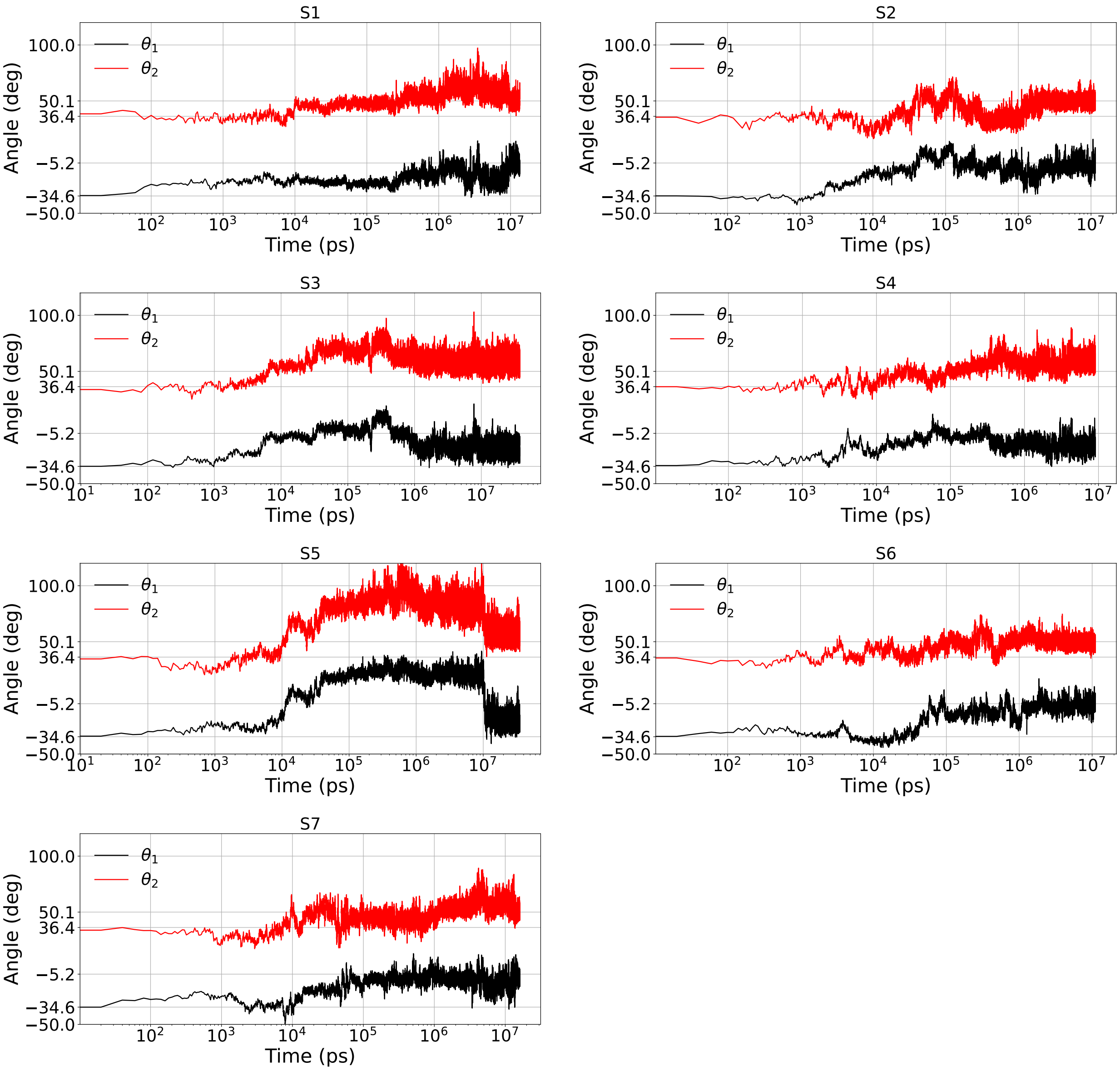


**Figure S5.** Time evolution of the two rotation angles $\theta_1$ and $\theta_2$ as defined in Figures S4 **B** and **C**, respectively, for all search simulations.The respective starting and average recognition angles at $-34.6°$ and $-5.2°$ for $\theta_1$, and at $36.4°$ and $50.1°$ for $\theta_2$ are indicated on the angle axis. Notice the logarithmic time scale.

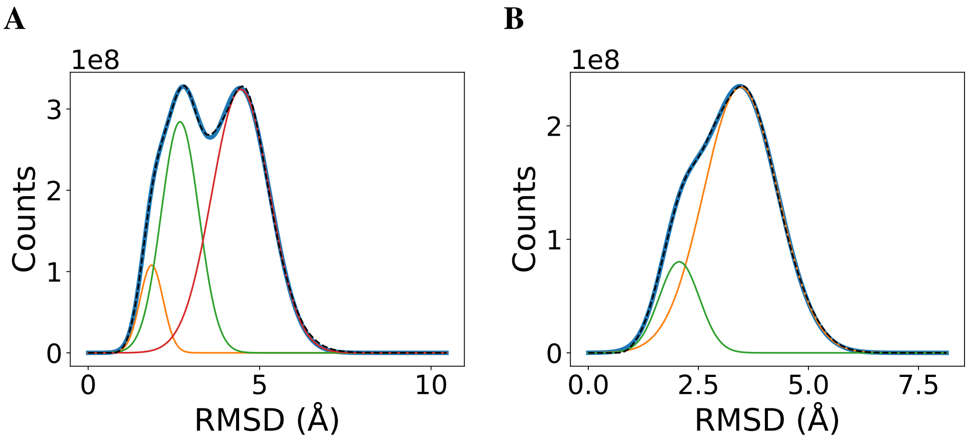


**Figure S6.** Distribution of pairwise RMSD values for **A** recognition, interrogation and search simulations combined, and **B** S189A mutation simulations shown as black dashed line. The fit is shown in blue as a sum of multiple Gaussian functions, which are also shown. The relevant parameters for **A** are $\mu = 1.85$ and $\sigma = 0.34$, $\mu = 2.68$ and $\sigma = 0.55$, and $\mu = 4.45$ and $\sigma = 0.84$ for the orange, green and red curve, respectively, and for **B** $\mu = 2.06$ and $\sigma = 0.46$, and $\mu = 3.46$ and $\sigma = 0.84$ for the green and orange curve, respectively.

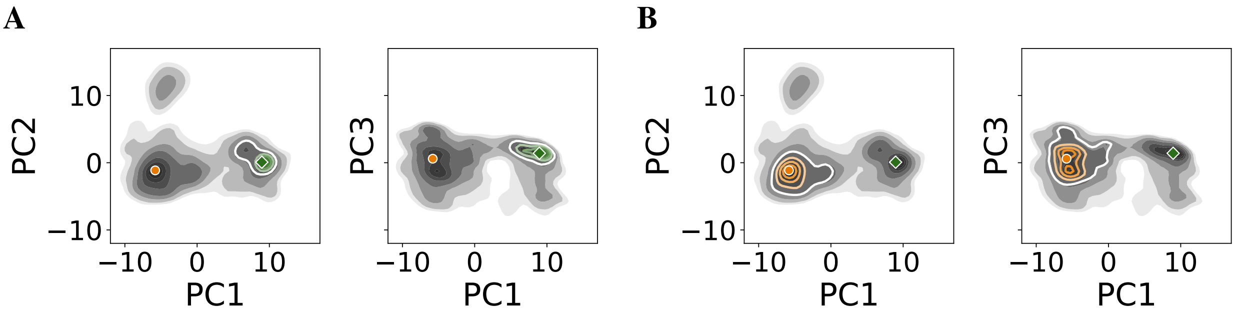


**Figure S7.** Projection of **A** cluster $C_1^{wt}$ and **B** cluster cluster $C_2^{wt}$ onto the first three principal components. The respective cluster centres are also highlighted as circle and diamond. The energy landscape of all data used for PCA is shown in grey colours.

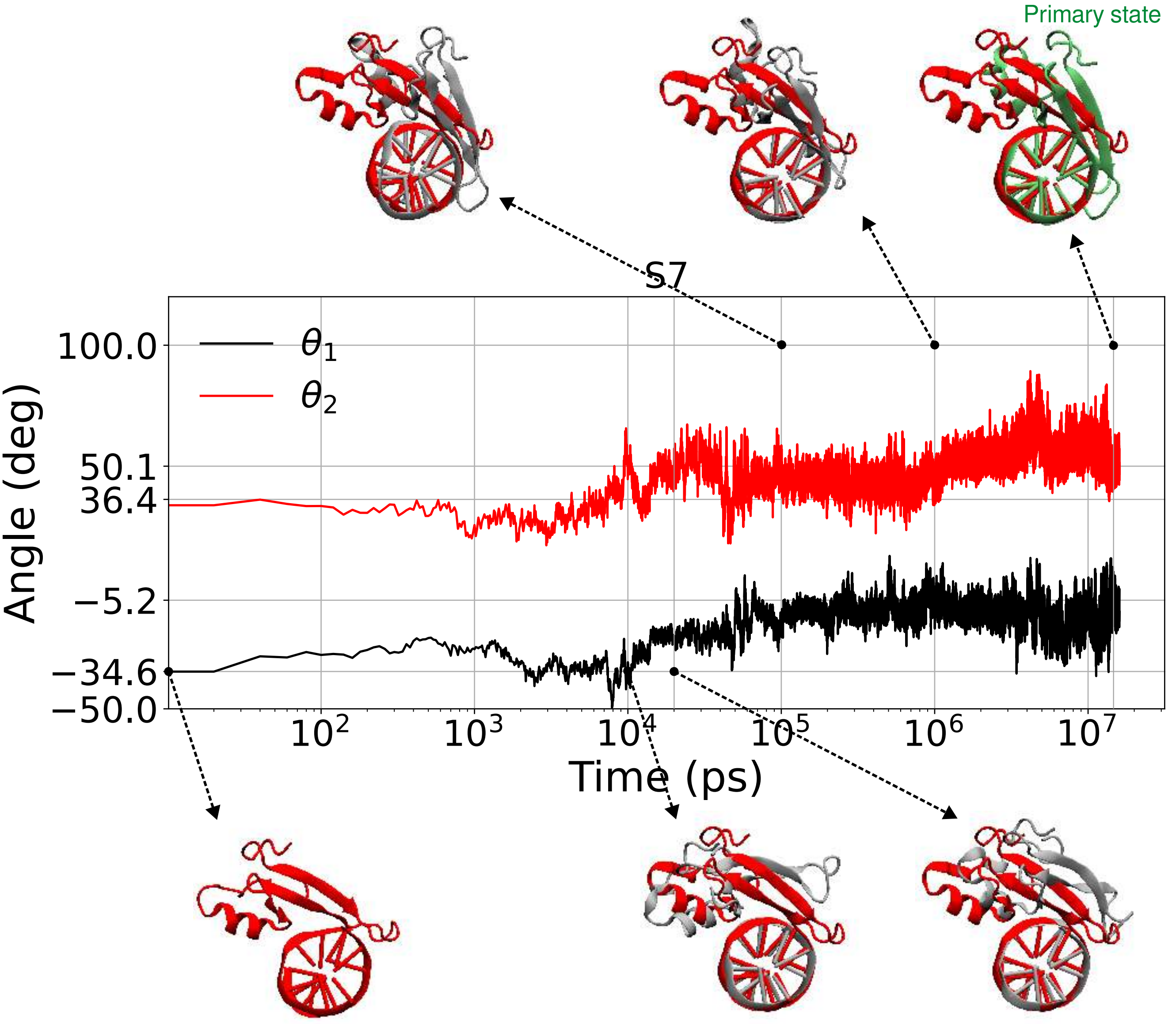


**Figure S8.** Time evolution of the orientational angles $\theta_1$ and $\theta_2$ during the search simulation (run S7), with representative snapshots of MBD2 positioning on DNA. The red structure shows the initial search configuration, in which MBD2 is positioned one base pair downstream of the specific mCpG site. Gray structures indicate conformations sampled at different simulation times, while the green structure corresponds to the primary bound state formed during the simulation. Together, the angular trajectories and snapshots capture the transition from a nonspecific search configuration toward a specific MBD2–mCpG conformation.

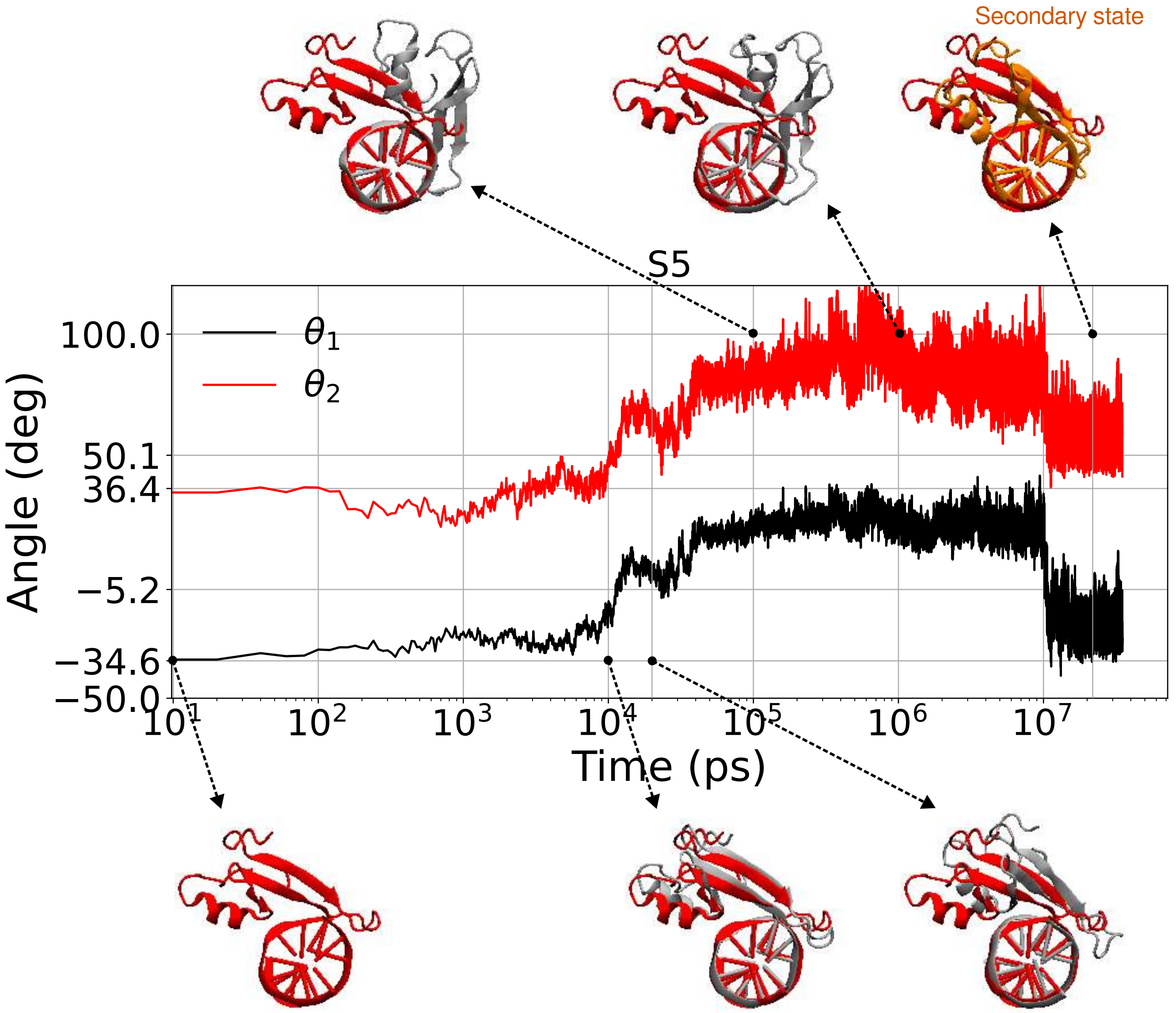


**Figure S9.** Time evolution of the orientational angles $\theta_1$ and $\theta_2$ during the search simulation (run S5), with representative snapshots of MBD2 positioning on DNA. The red structure shows the initial search configuration, in which MBD2 is positioned one base pair downstream of the specific mCpG site. Gray structures indicate conformations sampled at different simulation times, while the green structure corresponds to the primary bound state formed during the simulation. Together, the angular trajectories and snapshots capture the transition from a nonspecific search configuration toward a specific MBD2–mCpG conformation.

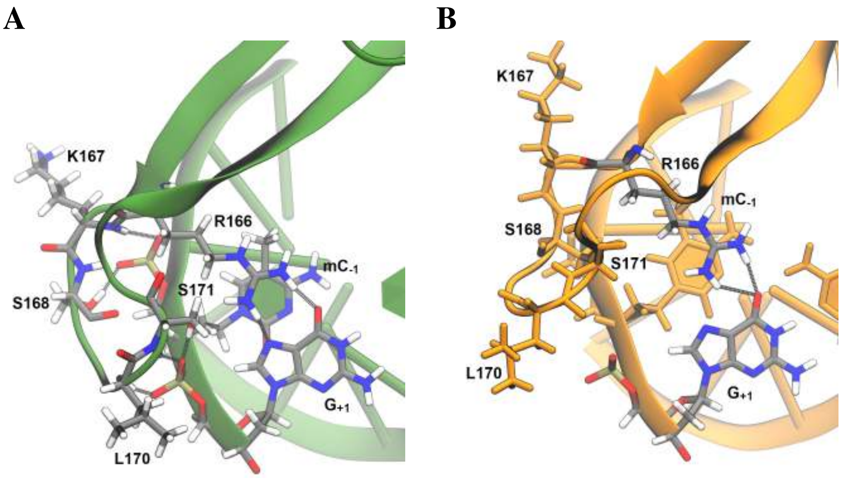


**Figure S10.** Selected detailed hydrogen bond interactions of the tail domain and the wild-type clusters **A** $C_1^{wt}$ and **B** $C_2^{wt}$. Residues involved in hydrogen bonds are coloured according to their atom type.

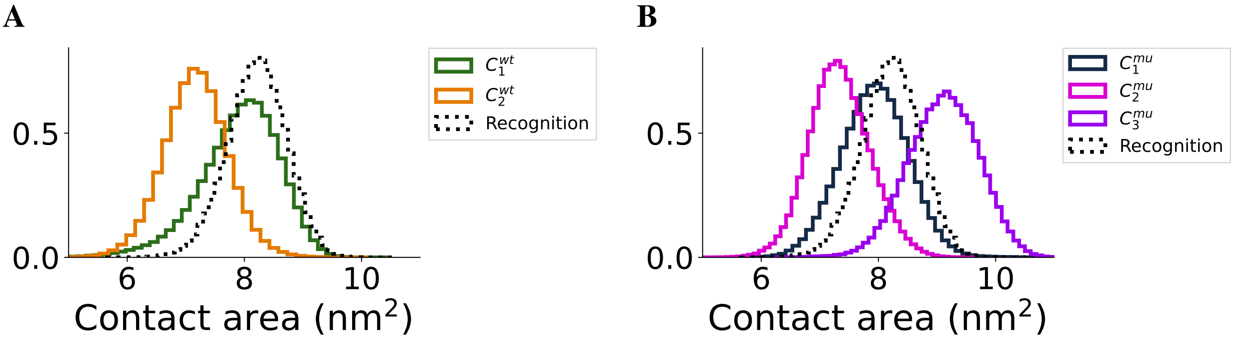


**Figure S11.** Contact area between protein and DNA for **A** the wild-type (WT) and the **B** S189A clustering using a threshold of $0.25$ nm and $0.22$ nm, respectively.

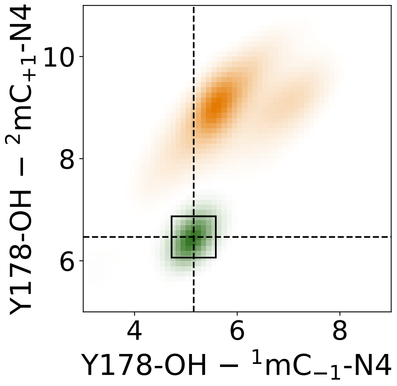


**Figure S12.** Distribution of the distances between the OH-atom of Y178 and the N4-atom of the methylated cytosines for the wild-type (WT) clusters. Dashed lines indicate the average values of the recognition simulations and the square is the standard deviation. Green and orange are clusters $C_1^{wt}$ and $C_1^{wt}$, respectively.

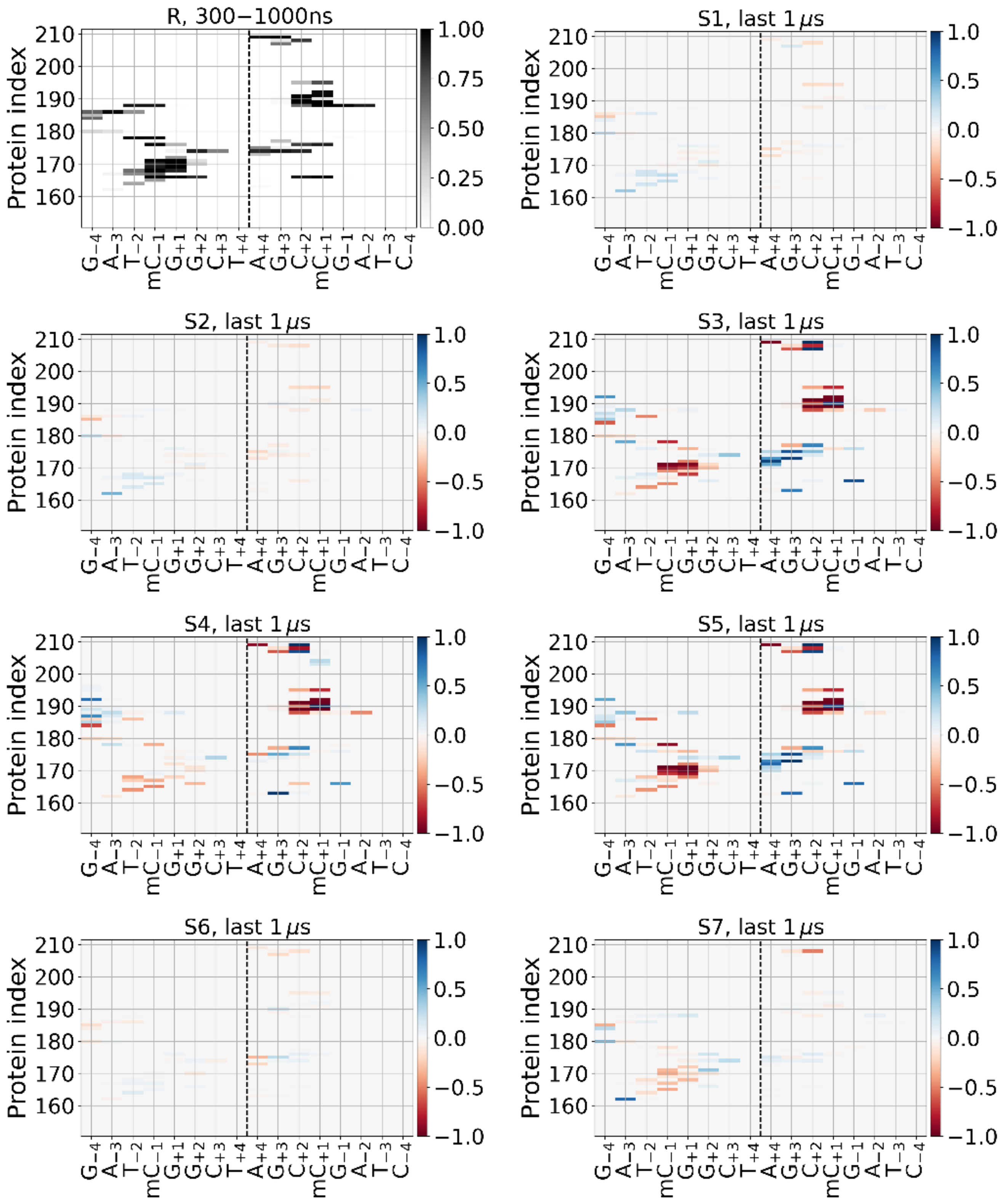


**Figure S13.** Contact maps of MBD2–DNA interactions are shown as the average over recognition simulation replicas (top left). For the last microsecond of each search simulation, differences relative to the recognition contact map are shown, with blue indicating stronger contacts and red indicating weaker contacts compared with the recognition simulations.

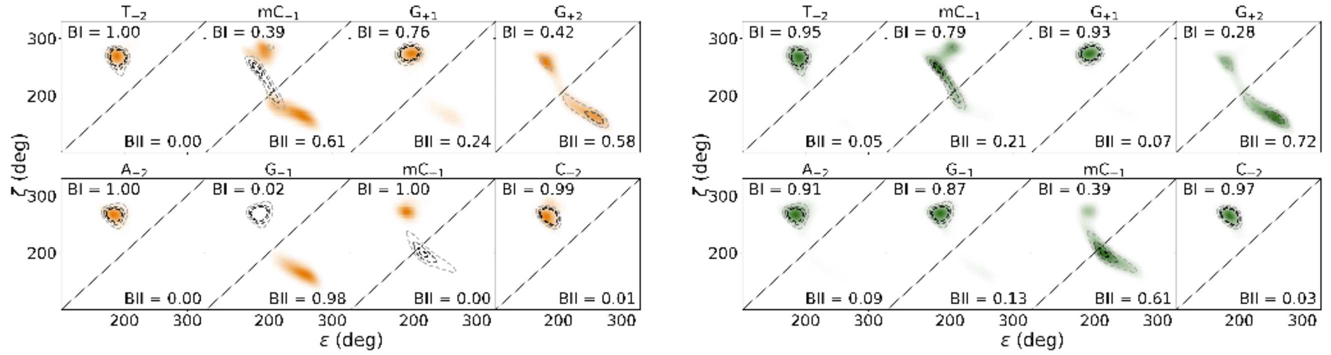


**Figure S14.** BI/BII conformation distribution for the central four basepairs, including the mCpG dinucleotide and the first two clusters $C_1^{wt}$ (green), $C_2^{wt}$ (orange), i.e. the primary and the secondary state, respectively. The fraction of the trajectory in BI and BII conformation is indicated in the respective area. The distribution for the recognition simulations is drawn as grey dashed lines.

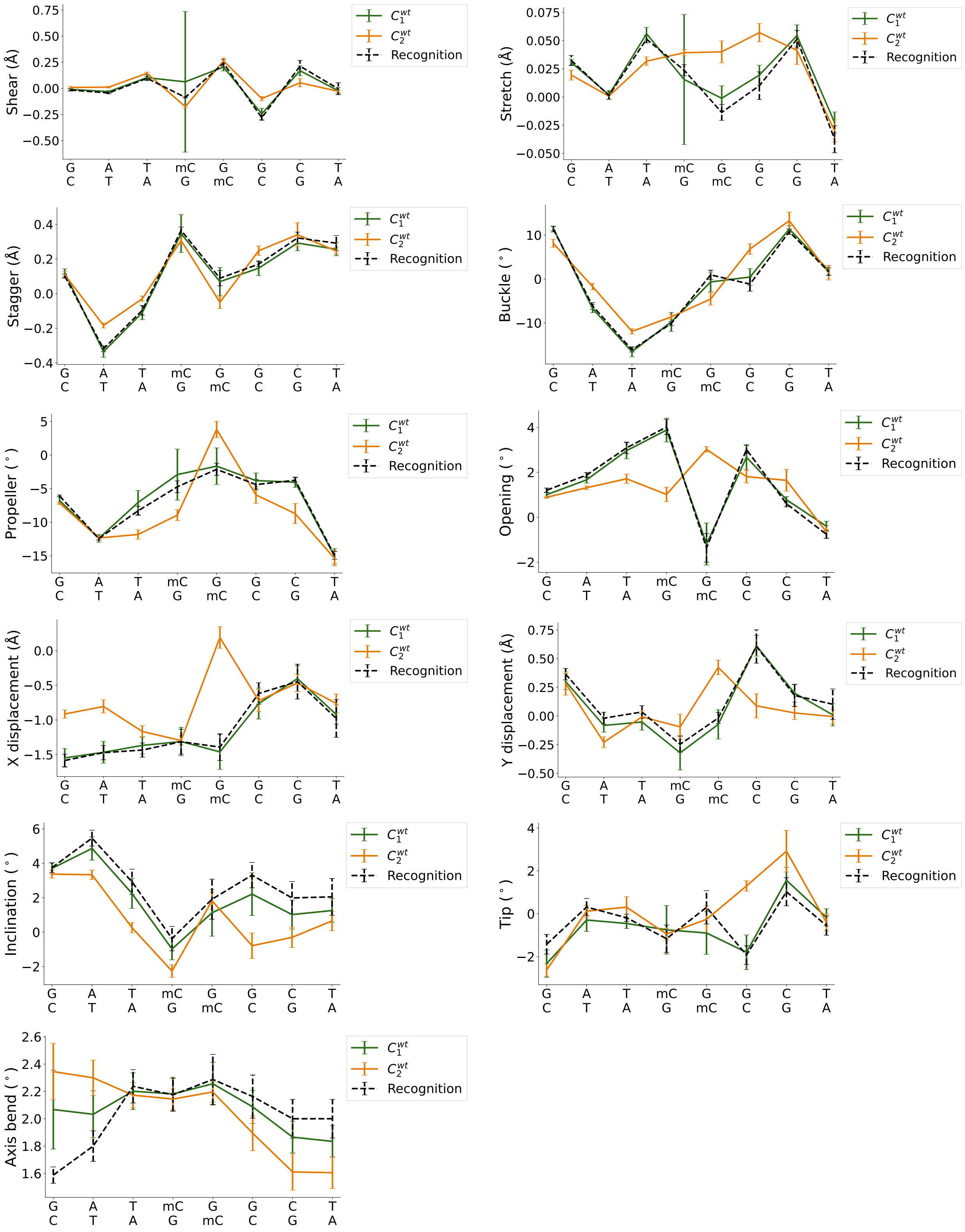


**Figure S15.** DNA base pair parameters for widtype (WT) clusters with a clustering threshold of 0.25 nm. The results for the recognition simulations are shown as reference.

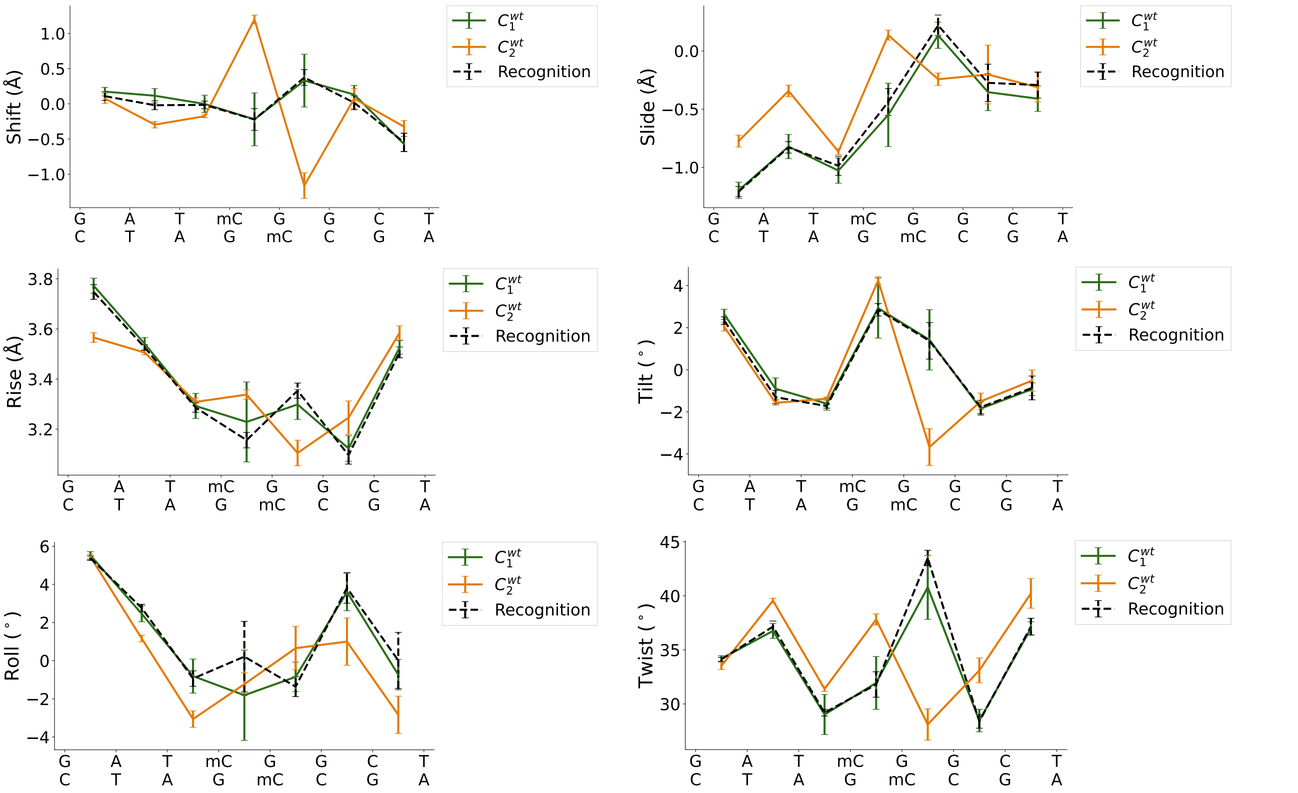


**Figure S16.** DNA base step parameters for widtype (WT) clusters with a clustering threshold of $0.25$ nm. The results for the recognition simulations are shown as reference.

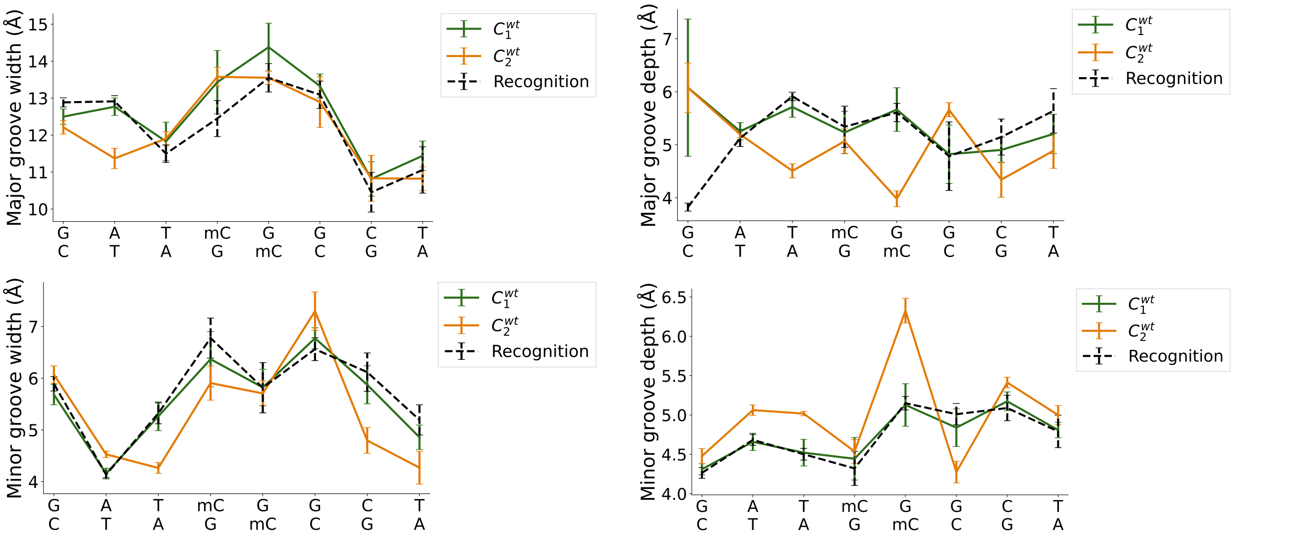


**Figure S17.** DNA major and minor groove parameters for widtype (WT) clusters with a clustering threshold of $0.25$ nm. The results for the recognition simulations are shown as reference.

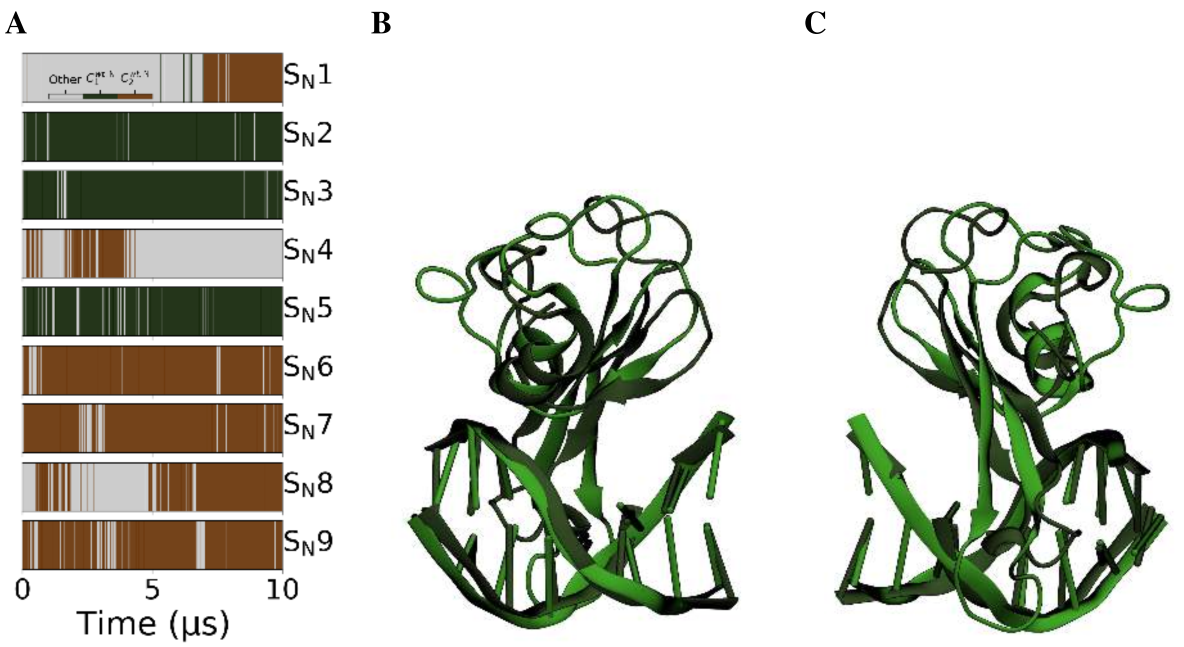


**Figure S18.** **A** RMSD clustering with a threshold of 2.5 Å of search simulations with a different DNA sequence (Figure 4C) also shows two main clusters $C_1^{wt,\mathrm{N}}$ (dark green), $C_2^{wt,\mathrm{N}}$ (brown). The distribution (not shown) has two main peaks separated by a valley similar to Figure S6**A**. Cluster center $C_1^{wt,\mathrm{N}}$ (dark green) and $C_1^{wt}$ (green) are shown as an overlay aligned to the DNA from **B** the front and **C** the back.

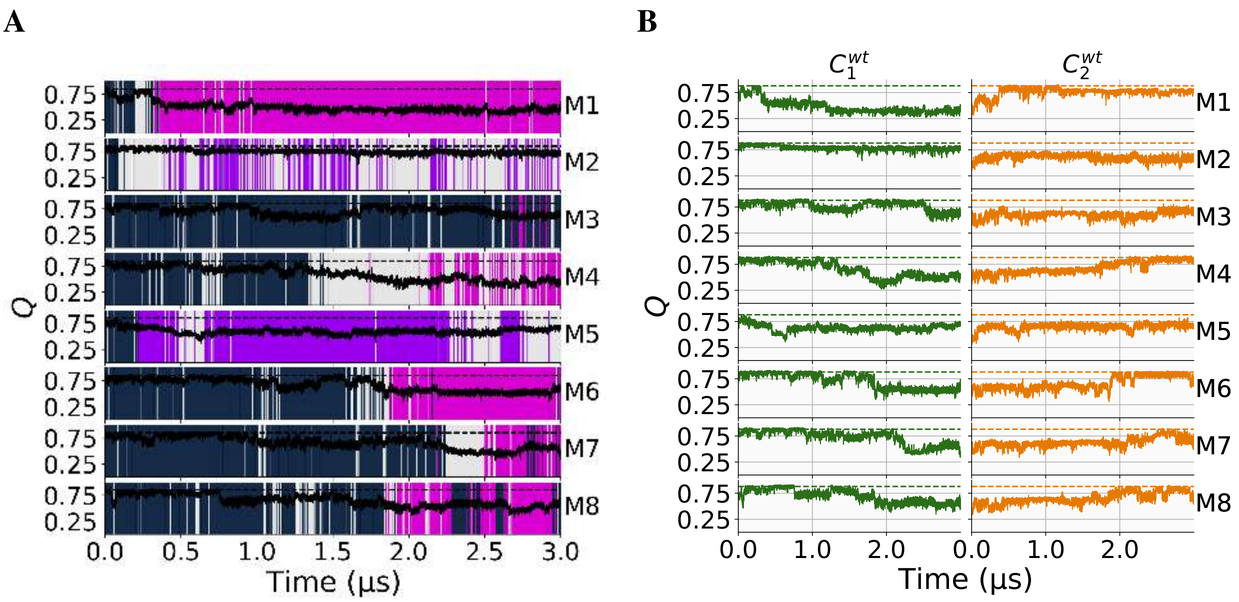


**Figure S19.** **A** RMSD clustering for the S189A mutation simulations with a threshold of 0.22 nm. The three clusters are $C_1^{mu}$ (blue), $C_2^{mu}$ (pink) and $C_3^{mu}$ (purple), respectively. The fraction of native contacts with respect to the reference model is shown in black with the average of the recognition simulations as a black dashed line. **B** Fraction of native contacts with respect to the cluster centers of the WT RMSD clustering. The average fraction of native contacts of the reference cluster is shown as a dashed line.

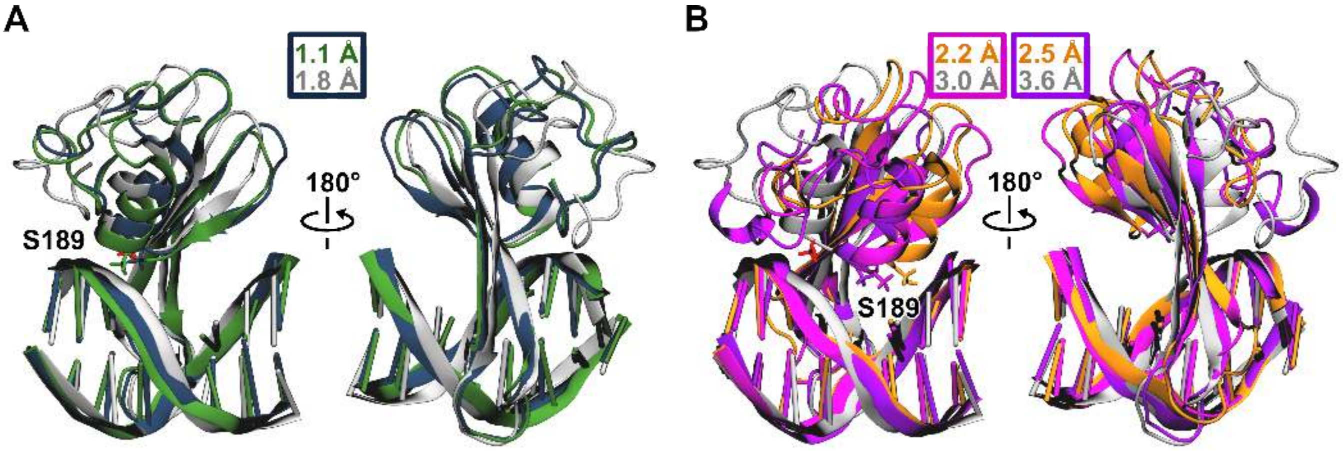


**Figure S20.** **A** Comparison of cluster center $C_1^{wt}$ (green) obtained from the wild-type simulations and the cluster center $C_1^{mu}$ (blue) obtained from the S189A mutation simulations. **B** Comparison of cluster center $C_2^{wt}$ (orange) obtained from the wild-type simulations and the second ($C_2^{mu}$, pink) and third ($C_3^{mu}$, purple) cluster center obtained from the S189A mutation simulations. For comparison, the reference model is shown in gray. The mutation is highlighted. RMSD values of the S189A mutation simulations cluster centers with respect to the other structures are printed in the respective colors.

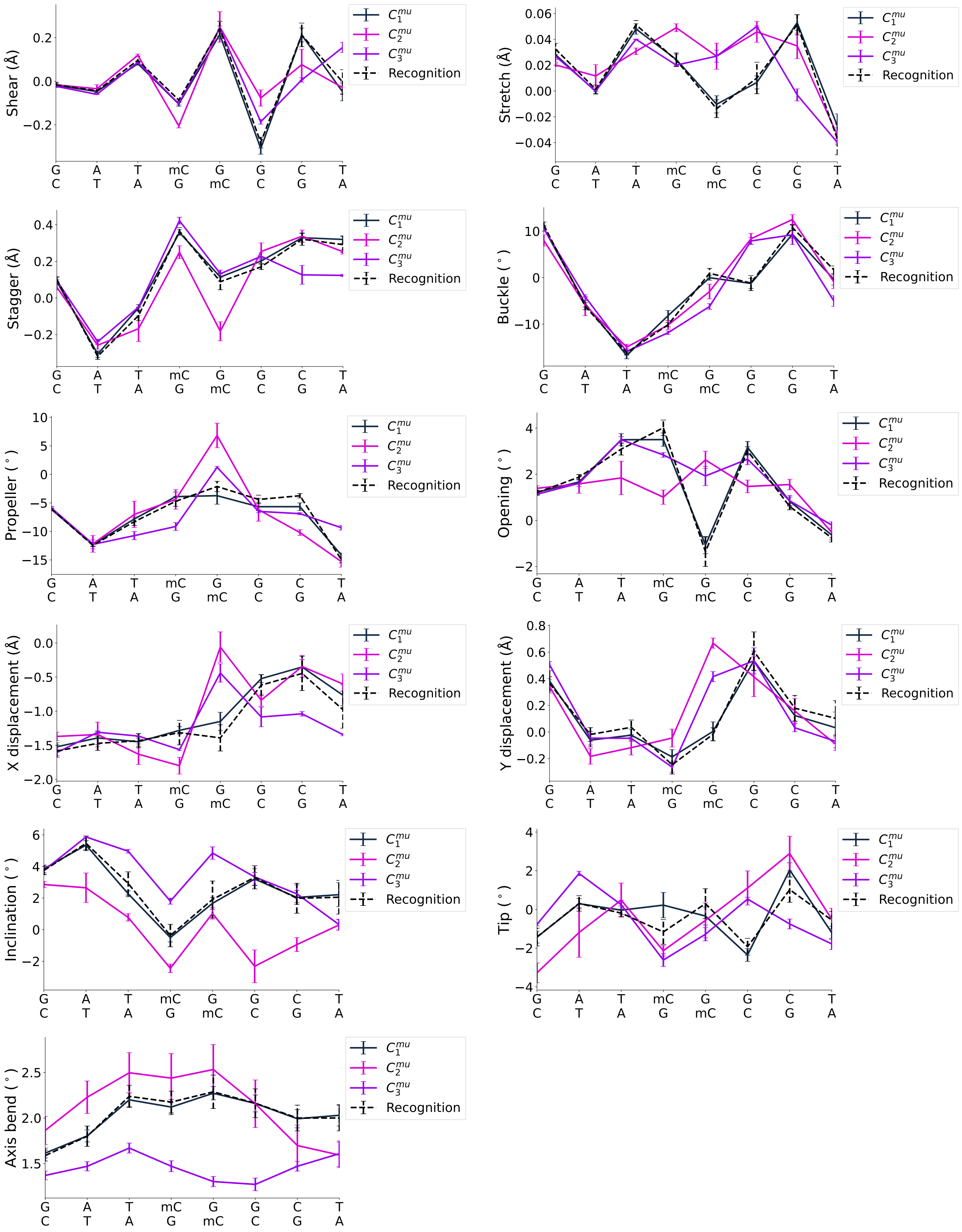


**Figure S21.** DNA base pair parameters for S189A clusters with a clustering threshold of $0.22$ nm. The results for the recognition simulations are shown as reference.

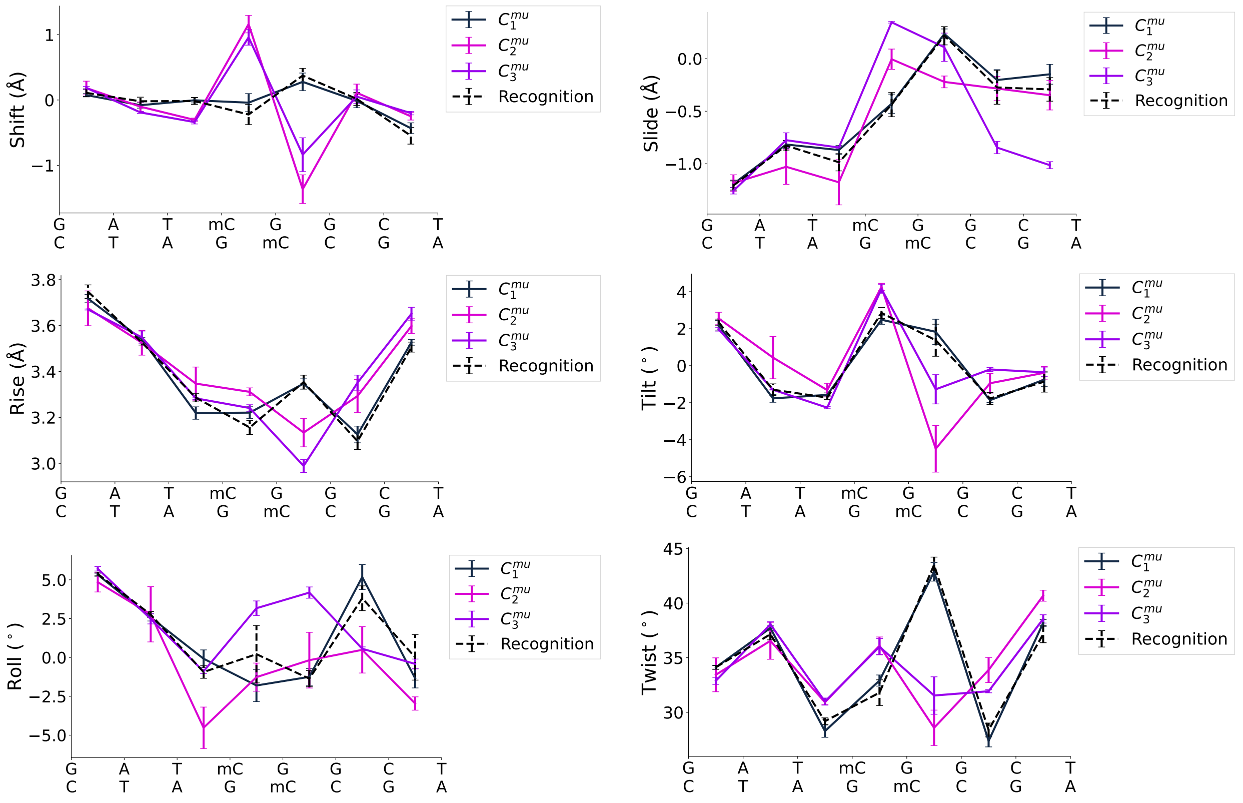

**Figure S22.** DNA base step parameters for S189A clusters with a clustering threshold of $0.22$ nm. The results for the recognition simulations are shown as reference.

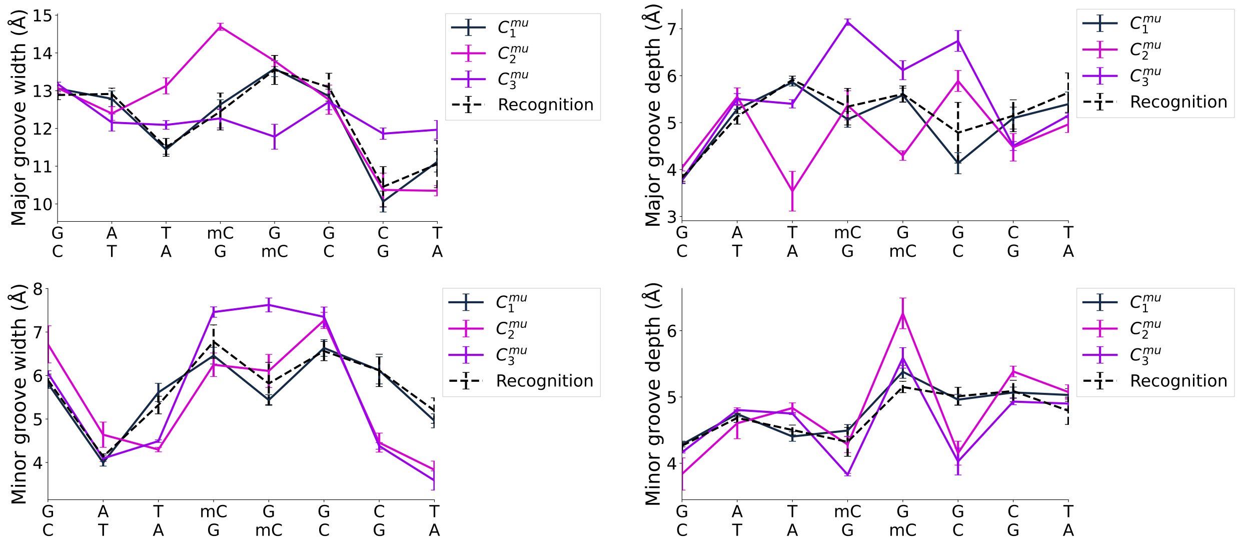

**Figure S23.** DNA major and minor groove parameters for S189A clusters with a clustering threshold of $0.22$ nm. The results for the recognition simulations are shown as reference.

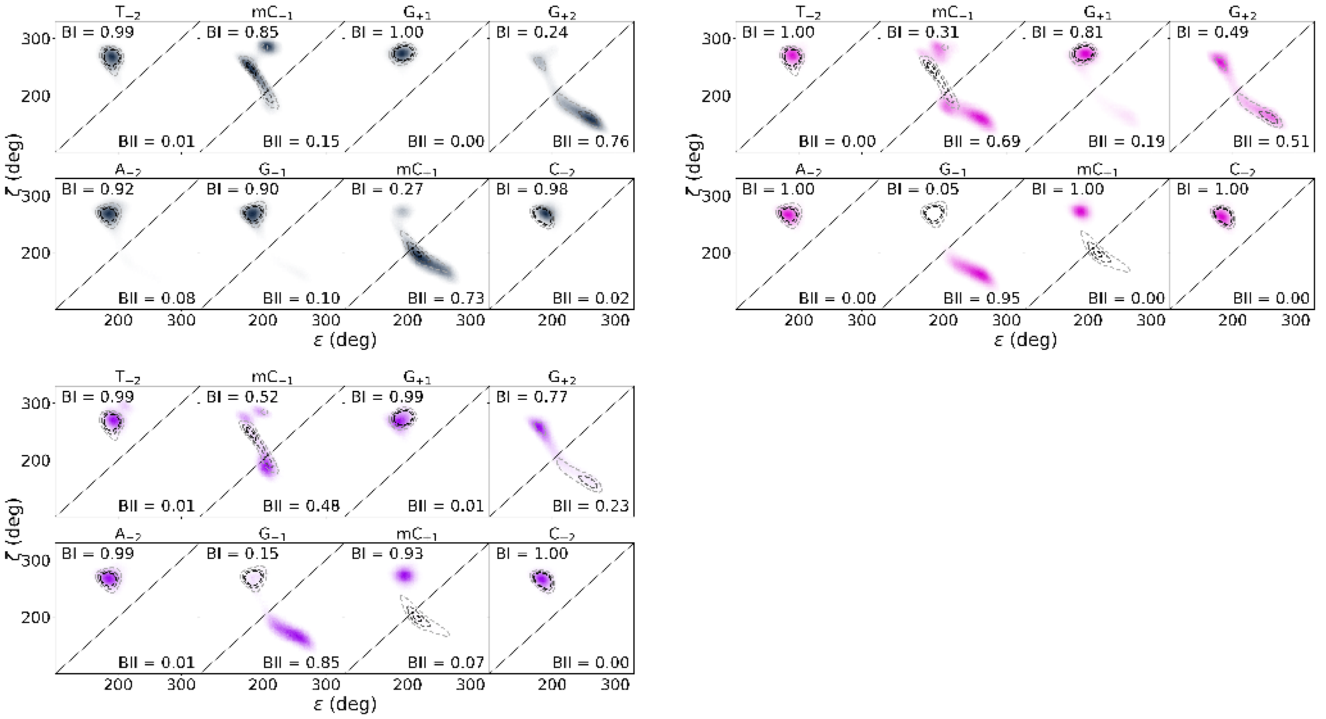


**Figure S24.** BI/BII conformation distribution for the central four basepairs, including the mCpG dinucleotide and the first three S189A clusters $C_1^{mu}$ (blue), $C_2^{mu}$ (pink), $C_3^{mu}$ (purple). The fraction of the trajectory in BI and BII conformation is indicated in the respective area. The distribution for the recognition simulations is drawn as gray dashed lines.

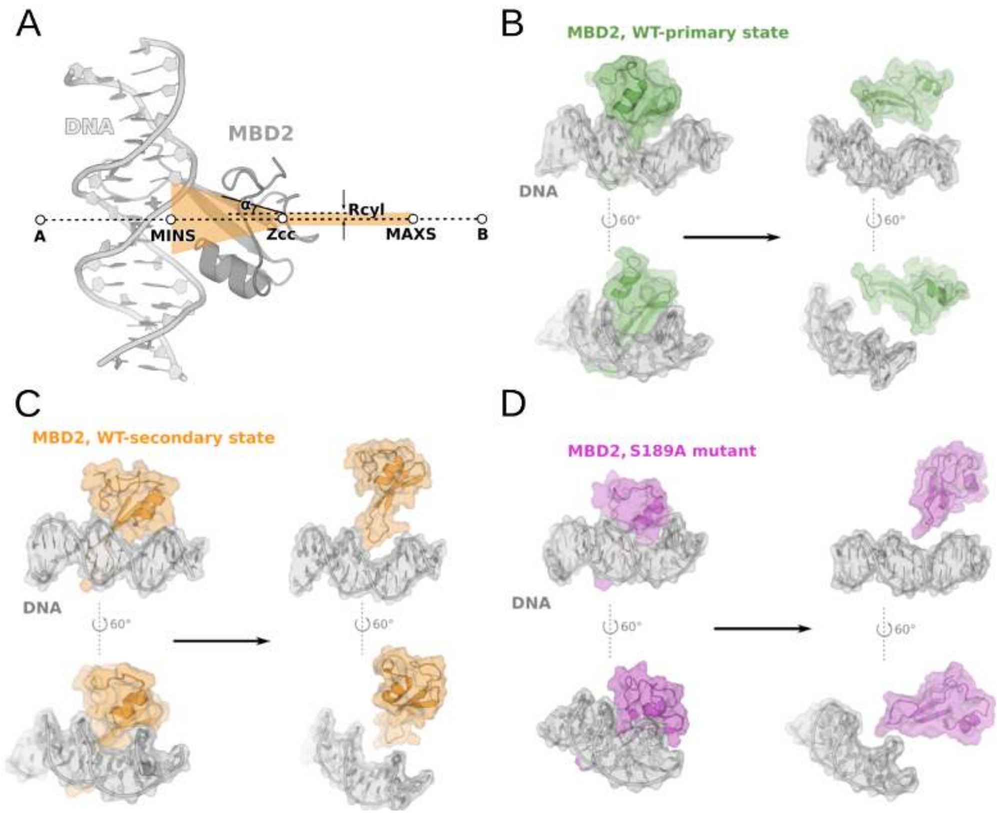


**Figure S25. A** Schematic representation of the funnel potential setup in PLUMED. Points A and B define the long axis of the funnel. In our setup, this axis passes through the centers of mass (COM) of the DNA and the MBD2 protein. The points labeled MINS and MAXS specify the boundaries (walls) that restrict the linear section of the funnel. The parameter Zcc marks the position where the cylindrical part of the funnel transitions into the conical (rotor) section. Rcyl defines the radius of the cylindrical region, while $\alpha$ specifies the angle of the conical section. **B–D** Bound and unbound states of the WT-primary, WT-secondary and S189A-mutant conformations. Front (upper) and side (lower) views.

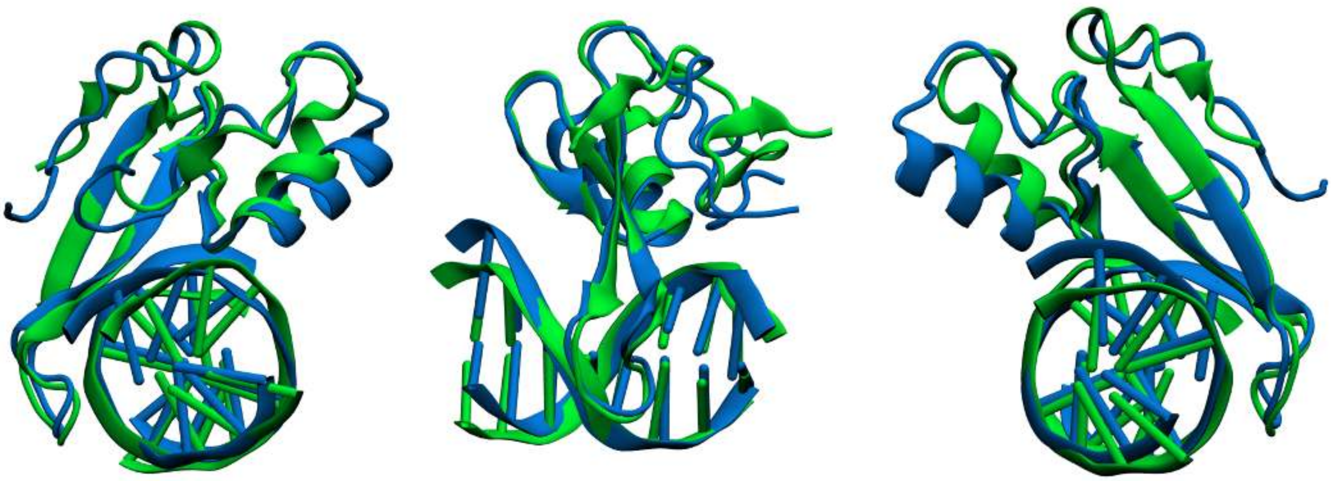

**Figure S26.** Overlay of the core regions of the MBD2–mCpG crystal structure with PDB ID: 7MWK (green) and the MeCP2–mCpG complex with PDB ID: 3C2I (blue), highlighting their similar protein–DNA conformations.

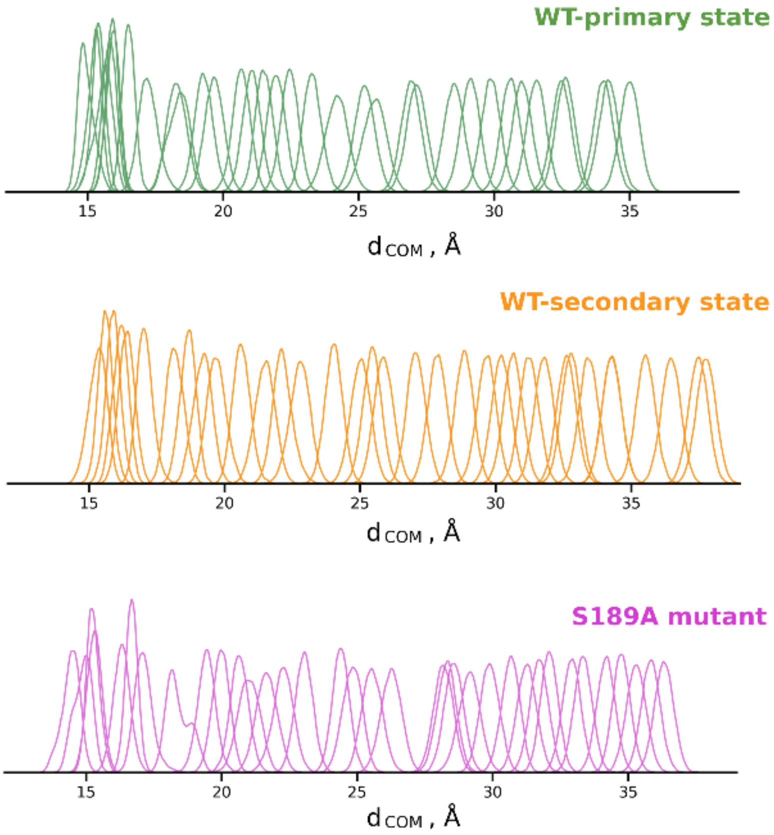


**Figure S27.** Umbrella Sampling windows (N = 35) covering the reaction coordinate (dCOM) in simulations of the WT-primary, WT-secondary, and S189A-mutant states of MBD2.

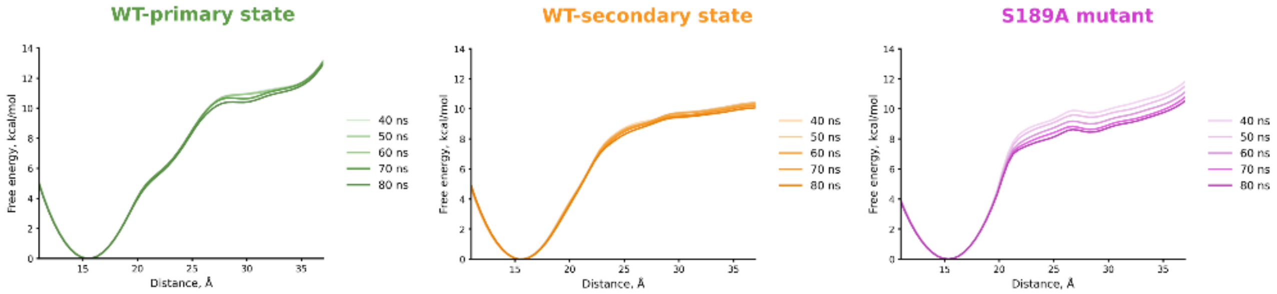


**Figure S28.** Convergence of the free energy profiles for the WT-primary, WT-secondary, and S189A MBD2-DNA systems.
